# Insight into the structure and interactions of the *M. tuberculosis* Mce-associated membrane proteins Mam1A-1D

**DOI:** 10.1101/2025.03.26.645568

**Authors:** Mikko J. Hynönen, Perumal Perumal, Nora Tir Hynönen, James J. Doutch, Kun Ma, Rajaram Venkatesan

## Abstract

Tuberculosis (TB) is a major infectious disease caused by *Mycobacterium tuberculosis* (Mtb), leading to more than a million human deaths every year. Mammalian cell entry complexes (Mce1-4) play an essential role in the survival of Mtb during the latent stage by mediating the import of lipids, including fatty acids and cholesterol, from the host. The proper functioning of Mce-complexes requires additional proteins such as Mce-associated membrane (Mam) proteins and lipid uptake coordinator (LucA), thus making them potential candidates for the development of anti-TB drugs. Four Mam (Mam1A-1D) proteins are coded from the *mce1* operon and two from the *mce3* (Mam3A-3B) and *mce4* (Mam4A-4B) operons. In addition, five orphaned mam (Omam) proteins have been identified, which are not part of the *mce* operons but are functionally relevant for the Mce complexes. Analysis of the sequences of Mam/Omam proteins suggests that they share many common secondary and tertiary structural elements despite the low sequence identity between them. Here, we have characterized a recombinantly produced Mam1A variant by small-angle X-ray and neutron scattering. The studies indicate that Mam1A is tetrameric in solution with two disulfide bridges necessary for the stability of Mam1A. Similarly, a disulfide bridge has also been identified in Mam1C. Furthermore, through coexpression and copurification, we demonstrate that Mam1A-1D and LucA interact to form stable Mam1ABCD as well as Mam1ABCD-LucA complexes. The results obtained pave the way for further understanding how the Mam1ABCD and Mam1ABCD-LucA complexes are organized and interact with the Mce complexes, leading to their stabilization.

## Introduction

Tuberculosis (TB) is one of the oldest diseases that has riddled mankind. Recent reports from the World Health Organization (WHO) estimated that nearly 11 million people fell ill with TB, leading to 1.4 million deaths [1]. The more recent reports also show that TB cases are rising everywhere except in the European region [2]. With the increase in multidrug-resistant strains detected and more strains resistant to conventional BCG vaccines, finding new ways to battle TB is crucial. *Mycobacterium tuberculosis* (*Mtb*), the pathogen that causes TB, has survived for centuries by adapting to the host intracellular environment. *Mtb* persists in its host in a latent form even for decades, during which *Mtb* switches from using carbohydrates to lipids as the main source of carbon and energy for its survival [3,4]. *Mtb* has an extensive repertoire of lipid-processing enzymes and protein complexes [5]. One such set of four homologous ATP-binding cassette (ABC) transporters is mammalian cell entry (Mce) complexes, Mce1-4, encoded by the *mce1*-*mce4* operons. The Mce1 complex encoded by the *mce1* operon is responsible mainly for *Mtb*’s palmitate and oleate intake [6]. Mutational studies have also suggested the role of the Mce1 complex in the import of mycolic acid [7]. Additionally, mutations of the *mce1* operon have been observed to cause hypervirulence [8]. The Mce4 complex is a cholesterol importer [9–11]. The Mce2 complex is likely a sulfolipid transporter [12]. The precise substrate specificity of the Mce3 complex is not yet known. The importance of the Mce proteins for virulence [13–15] and survival of mycobacteria [14–16] has been shown in animal models.

The *mce1*, *mce3,* and *mce4* operons also code for additional proteins, which are named Mce-associated membrane (Mam) proteins [17]. The *mce1* operon codes for four Mam proteins (Mam1A-1D), while the *mce3* and *mce4* operons code for two Mam proteins each (Mam3A-3B and Mam4A-4B) [17]. Additionally, five proteins homologous to Mam proteins have been identified as encoded outside the *mce* operons, and these are named orphaned Mam (Omam) proteins [17,18]. *Mtb* has five *omam* genes, labelled *omamA*, *B*, *C*, *D*, and *E*. The Mam and Omam proteins are the least understood proteins among the Mce-related proteins. Although their importance in infection has been established with mutational studies [16,19], only recently, it has been suggested that OmamA plays a crucial role in stabilizing the Mce1 complex and enabling the Mce4 complex to work properly [18,20]. Moreover, it has been demonstrated that the Mam proteins are not involved in the binding of cholesterol to the Mce4 complex. Rather, they are involved in the internalization of cholesterol at the cytoplasmic membrane. The process involves another protein named LucA which interacts with the Mam/Omam proteins and is involved in the stabilization of the Mce1 and Mce4 complexes [6]. Deletion of LucA showed an increase in the expression of all *mce1* genes while almost completely inhibiting the palmitate lipid uptake [6,20]. Inhibiting the interaction of the Mam/Omam proteins with LucA and with the Mce complexes could be an effective strategy to impair the function of the Mce complexes, potentially reducing the survival of *Mtb* in its latent form. Therefore, a detailed understanding of the Mam/Omam proteins and LucA is necessary. In this study, we recombinantly produced Mam1 proteins, and through biophysical characterization, including SAXS and SANS studies, showed that Mam1A most likely exists as a tetramer. In addition, we showed that Mam1A-1D and LucA interact to form stable Mam1ABCD and Mam1ABCD-LucA complexes. Furthermore, we have discovered two crucial disulfide bonds in Mam1A and Mam1C that are important for their overall stability. These studies pave the way for further detailed structural and interaction studies on Mam1ABCD and Mam1ABCD-LucA complexes.

## Results

### Sequence analysis of the Mam/Omam proteins

Analysis of the sequences of Mam1A-1D suggests that Mam1A and Mam1C share a sequence identity of 21%, whereas Mam1A and Mam1B have only about 4% sequence identity (**Supplementary Table 1**). Despite the low sequence identities between them, secondary structure predictions suggest that Mam1A-1D have a common domain organization with four key regions (**Fig. 1A and Supplementary Figure 1A**). The N-terminal region has a varying number of amino acids; for example, 28 amino acids in Mam1C is the shortest, and 159 in Mam1B is the longest region. The Mam1B variable region additionally contains many small α-helices and β-strands, which do not occur in the other Mam1 proteins. The variable N-terminal region is followed by a long α-helix that contains a transmembrane (TM) region. While Mam1A, Mam1C and Mam1D have only one predicted TM helix, Mam1B is predicted to have three TM helices, two of which are in the variable region (**Supplementary Figure 2**). In addition, Mam1B has one more helix predicted to be buried in the membrane (**Fig. 1A**). The TM region in each Mam is about 20 amino acids long, and the α-helix containing TM is the longest in all Mam1 proteins, ranging from 43 to 55 amino acids. After the α-helix-containing TM region is a small helical region with two α-helices, each of which is between 10 and 20 amino acids long. The protein C-terminus contains a β-sheet segment with four β-strands. The β-strands are between 7 and 14 amino acids long.

**Fig. 1.**
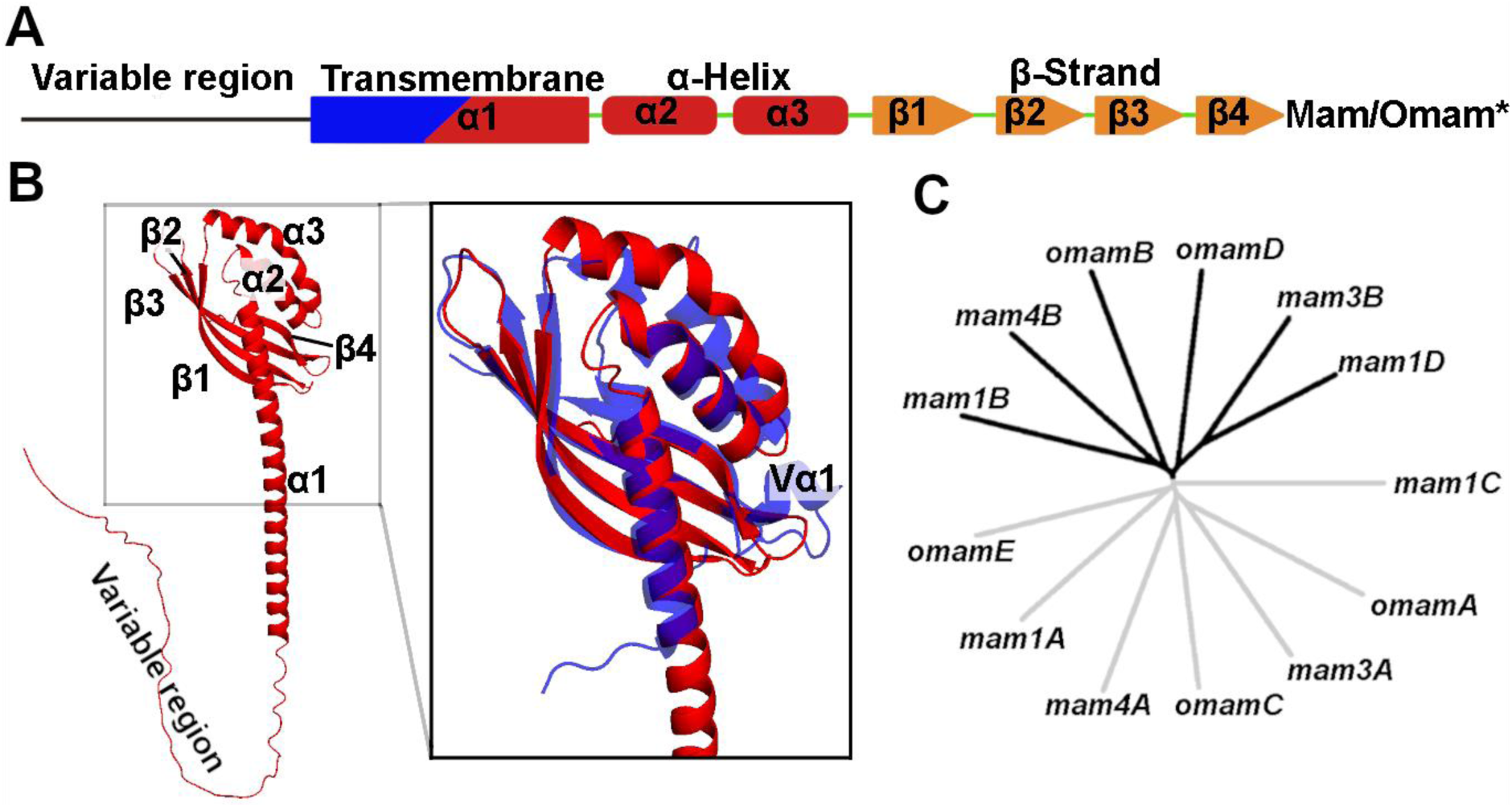
Sequence analysis and secondary and tertiary structure predictions of Mam and Omam proteins. **(A)** A schematic illustration of the general conserved secondary structure organization of all *Mtb* Mam/Omam proteins excluding Mam1B and OmamB (denoted as Mam/Omam*). The four regions are named i) variable region, ii) transmembrane iii) α-helix and iv) β-strand. **(B)** AlphaFold2-predicted tertiary structure of Mam1A in cartoon representation (red), a representative of a general Mam/Omam protein. The prediction quality parameters are summarized in **Supplementary Table 3**. Magnified view of C-terminal domains of Mam1A superimposed onto the crystal structure of VirB8 (PDB id: 2cc3 chain A, blue cartoon). RMSD score of superimposed models is 1.325 for all Cα atoms. VirB8 structural elements line up with Mam1A, except for VirB8 having an extra helix between β3- and β4-strands, annotated as Vα1. **(C)** Phylogenetic tree generated from the sequence alignment of all Mam/Omam proteins of *Mtb,* highlighting that Mam/Omam As, Cs and E cluster on one side of the tree and Mam/Omam Bs and Ds cluster together on the other side of the tree, dividing the tree into two halves.

Sequence comparison shows that there are only four amino acids that are conserved in all Mam1A-1D (**Supplementary Figure 1**). Two of these, arginine (Mam1A-R52, Mam1B-R162, Mam1C-R25, Mam1D-R77) and alanine (Mam1A-A61, Mam1B-A170, Mam1C-A34, Mam1D-A86), are located close to or inside the α1-helix (**Fig. 1B**). The α1-helix is the longest helix in Mam1 proteins, which also contains a TM region in the first half of the helix. The α3-helix contains a conserved glutamine (Mam1A-Q131, Mam1B-Q237, Mam1C-Q103, Mam1D-Q154) and the β1-strand a conserved valine (Mam1A-V159, Mam1B-V262, Mam1C-V130, Mam1D-182) (**Supplementary Figure 1**). The importance of these conserved amino acids for the potential function of the proteins is not yet known. Mam1B has been noted to contain a conserved RDD domain. This is part of the RDD superfamily, characterised by an arginine and two aspartates as well as two TM regions [21]. This superfamily’s function is unknown, and no other proteins within the family have been studied. The three RDD conserved amino acids in Mam1B are R95, D133 and D144, which are absent in other Mam and Omam proteins. The proposed RDD domain is highlighted in **Supplementary Figure 2B** with a black box. Overall, sequence-related properties of each Mam and Omam are listed in **Supplementary Table 2**.

Mam proteins from the *mce3* and *mce4* operons and OmamA-E have a similar secondary structure organization. The variable N-terminal regions for Mam3B and Mam4B are much shorter, with less than 6 amino acids, immediately followed by the TM region (**Fig. 1A**). Additionally, OmamB has three TM regions similar to Mam1B. However, OmamB does not have as long a variable region as in Mam1B and the helix is predicted to be buried in the membrane (**Supplementary Figure 2C**). Comparison of all the Mam/Omam sequences shows that overall, Mam/OmamAs/Cs and OmamE fall under one branch of the phylogenetic tree while Mam/OmamBs/Ds fall under another branch (**Fig. 1C**). Between Mam/Omam proteins, the sequence similarity is much higher at the C-terminus of the proteins, where the alignment of the secondary structure elements is better. Overall, the highest identity is 31% between Mam3B and OmamD, and the lowest is between Mam1A and Mam1B (**Supplementary Table 1**). The TMHMM2.0 [22] prediction provides information on the TM region as well as the overall topology. The prediction confidence for Mam/Omam orientation varies between proteins (**Supplementary Table 2**).

A comparison of the AlphaFold2[23] predicted structures of each of the Mam/Omam proteins showed that they have many common features and are generally in line with the secondary structure predictions (**Fig. 1A and 1B**). In the predicted structures, the β-sheet partially surrounds the backbone α1-helix and the following α2 and α3 helices to form a V-shaped hinge, indicating flexibility in the structure. The β-sheet with the α2 and α3 helices creates a pocket for possible interactions. It has been shown that OmamA has a similar predicted structure as VirB8 using a Phyre 2 web service [18] (**Fig. 1B**). Like Mam and Omam proteins, VirB8 is a membrane protein with a single TM region. The structure of VirB8 has been determined by crystallography [24]. The similarities between VirB8 and Mam/Omam proteins are threefold: i) they have one long α-helix that is partially surrounded by the β-sheets at the C-terminus, which also contains the TM region, ii) the longer α-helix is followed by two shorter α-helices that form a V-shape turn, and iii) the protein ends in a β-strand-rich region with short loops connecting each. VirB8 has an additional small helix (Vα1) in the region connecting the β-stands β3 and β4. Comparing the Mam1A predicted C-terminal domain structure to the crystal structure of VirB8 (PDB ID 2cc3, chain A) yields a root mean square deviation (RMSD) of 1.325 for Cα atoms. However, it should be noted that overall, the AlphaFold parameters for the individual members of the Mam and Omam family are relatively poor (**Supplementary Table 3**).

### Mam1A_107-213_ forms a homotetramer with an intermolecular disulfide bridge

All the Mam/Omam proteins have a predicted transmembrane helix in the N-terminus. To increase the solubility, a variant of Mam1A was created by removing 106 residues from the N-terminus (Mam1A_107-213_), expressing only the C-terminal region outside the membrane. Preliminary expression and purification indicated that Mam1A_107-213_ still required the presence of detergents, similar to the full-length Mam1A. However, Mam1A_107-213_ required detergent only in the lysis step of the purification for obtaining a well-folded homogenous protein sample (**Fig. 2A**). Dodecyl nonaethylene glycol ether (C12E9) was chosen as the most optimal detergent for further purification as the use of this detergent for purification resulted in the lowest polydispersity measured by the dynamic light scattering (DLS) (**Supplementary Figure 3**) and the best unfolding curve, as observed by nano differential scanning fluorimetry (NanoDSF) suggesting a well-folded protein (**Fig. 2B**). The identity of the purified protein was confirmed by peptide mass fingerprinting using matrix-assisted laser desorption/ionization time-of-flight mass spectrometry (MALDI-TOF MS).

**Fig. 2.**
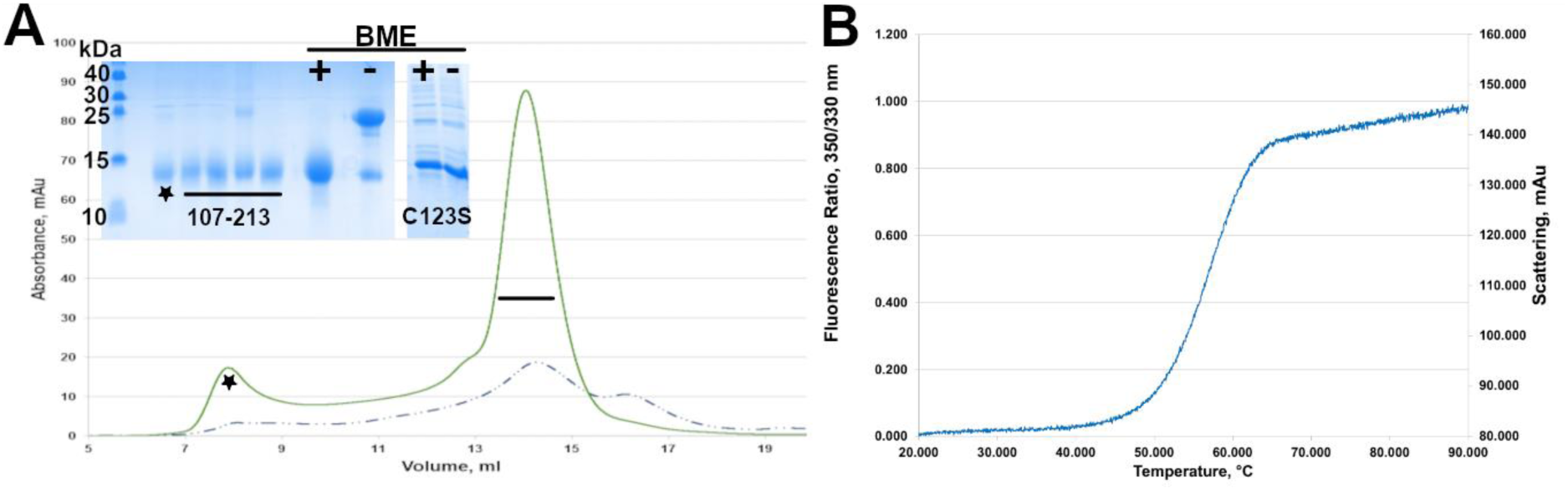
Purification profile and NanoDSF curve of Mam1A_107-213_. **(A)** SEC profile of Mam1A_107-213_(green line) comparison with Mam1A_107-213_C134S (dotted blue line). SDS-page insert highlighting the selected fractions for analysis with (+) and without (-) BME. Comparing the effect of BME in Mam1A_107-213_C134S shows that the two samples run identically. **(B)** NanoDSF profile of the purified Mam1A_107-213_ from the peak fractions (13.5-15 ml) from panel A.

Mam1A_107-213_ eluted at a retention volume of 13.5 ml in a Superdex 200 10/30 GL column (GE Healthcare), much earlier than expected for a protein with a theoretical monomeric molecular mass of 14.8 kDa (**Fig. 2A**). In the SDS-PAGE analysis the band corresponding to Mam1A_107-213_, as confirmed by peptide mass fingerprinting analysis, is observed at a higher molecular mass (∼30 kDa) close to that of a dimer in the absence of beta-mercaptoethanol (BME). In contrast, in the presence of BME, the band is observed close to the theoretical monomeric molecular weight (Fig. 2A). The primary sequence of Mam1A_107-213_ has only one cysteine (Cys123), which should, therefore, be involved in the disulfide bridge formation, stabilizing the homodimer. To test this, a variant Mam1A_107-213_-C123S was generated by mutating Cys123 to Ser. For this variant indeed, in SDS-PAGE, the band corresponding to Mam1A_107-213_-C123S is observed close to the theoretical monomeric molecular weight both in the presence and absence of BME, further confirming the involvement of Cys123 in disulfide bridge formation. Moreover, the Mam1A_107-213_-C123S variant was less stable and was prone to severe aggregation during concentration, suggesting the importance of this disulfide bridge for the overall stability of Mam1A_107-213_ (**Fig. 2A**). The yield and stability of the Mam1A_107-213_-C123S variant were too poor for further characterization.

The NanoDSF data was analyzed with MoltenProt [25], which indicated a melting temperature (*T*_m_) of 56.5 °C (**Fig. 3B and Supplementary Figure 4**). Analysis of the molecular mass of Mam1A_107-213_ by size-exclusion chromatography (SEC)- multi-angle light scattering (MALS) followed by protein-conjugate analysis showed that the total molecular mass of the eluted Mam1A_107-213_ was around 68 kDa, of which 54 kDa was protein and 14 kDa was C12E9 detergent. The measurement revealed that although the detergent was used only in the first lysis step, it was strongly bound to Mam1A_107-213_ (**Supplementary Figure 5**). Furthermore, these results suggest that the oligomeric state of Mam1A_107-213_ is tetrameric. Additional hydrodynamic radius calculation conducted by DLS measurement of the SEC-MALS run gave a radius of 3.45 nm, which corresponds to the combined mass of Mam1A_107-213_ tetramer surrounded by a detergent micelle of around 72 kDa. Therefore, the tetramer is proposed to contain two inter-subunit disulfide bridges formed between the two pairs of Cys123.

**Fig. 3.**
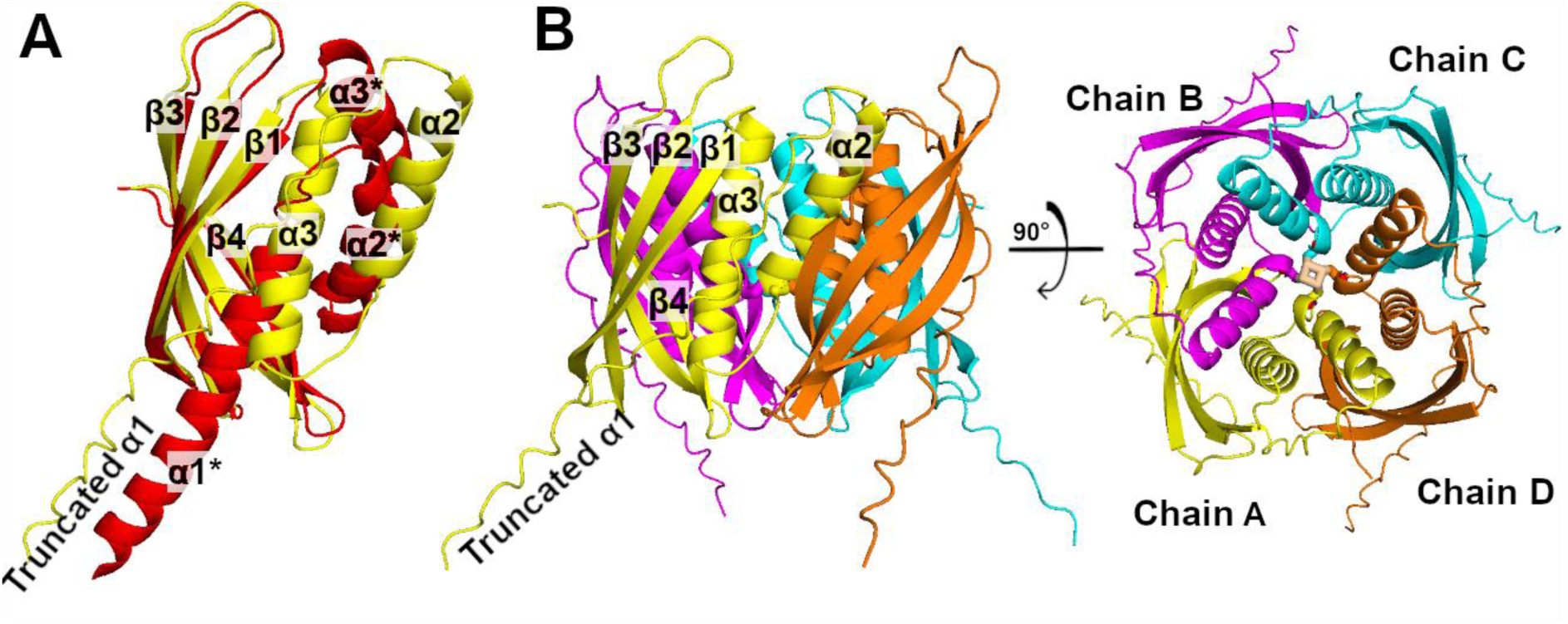
Predicted tetramer assembly of Mam1A_107-213_. **(A)** Predicted structure of Mam1A full length (red cartoon), superimposed to chain A of predicted tetrameric assembly of Mam1A_107-213_ (yellow cartoon). Secondary structure elements annotated with an asterisk belong to the full-length Mam1A prediction. **(B)** Selected tetramer model of Mam1A_107-213_. The tetramer model was selected from the predicted models as it satisfied the condition of placing cysteines close enough to form disulfide bonds, shown as sticks in the center of the predicted structure (wheat-colored sticks). Extending the chain is a predicted linker of the expression vector. It can be proposed that the α1 helix would occupy this region in the full-length model.

### Disulfide bridge formation is possible only by helix swapping

The closest known homolog of Mam1_107-213_, VirB8, forms a dimer in the crystal structure [24], whereas Mam1A_107-213_ forms a tetramer. Superposition of the predicted monomeric Mam1A_107-213_ structure on the VirB8 dimers did not bring the two Cys123 residues close enough to form a disulfide bridge. Mam1A_107-213_ did not crystallize despite multiple trials. Therefore, we predicted the tetrameric assembly of the variant using AlphaFold3[26]. All the predicted tetramers showed very low confidence global fold accuracy (pTM) of 0.1 to 0.25 and high predicted aligned error (PAE) of 15-28 Å. Per chain pTM is slightly higher, 0.41, but still at low confidence. Low scoring could also be the result of having no close representative models of such symmetry. It should be noted that the pTM for the full-length Mam1A is only 0.62. Local confidence parameters (pLDDT) between full-length and truncated chains differ such that in full-length Mam1A, β-strands have pLDDT values above 90, while in the truncated variant, the strands have values between 70 and 90. The α2- and α3-helices have pLDDT values of 70 to 90 in full-length Mam1A prediction, while in truncated, the values are between 50 and 70. The global pLDDT for full-length Mam1A prediction is 75.32, while for truncated Mam1A chain in the proposed tetramer model, the global pLDDT is 38.73. Prediction information for other Mam and Omam proteins is given in **Supplementary Table 3**.

Further analysis of the predicted models revealed two candidate models in which the Cys123s are close enough to form disulfide bridges and do not have clashes between the chains. The final selection of the complex was chosen based on the placement of Cys123, as well as the fit to the dummy atom model of SAXS and SANS. Superposition of the individual subunits from the selected tetramer model to the predicted monomeric structure of Mam1A showed that while the β-strands superpose well with each other, the α2 and α3 helices are placed differently when compared to the predicted monomeric Mam1A (**Fig 3A**). All the 4 chains have similar conformation in the tetramer model leading to a symmetrical tetramer (**Fig. 3B**). In this model, the cysteines are located so that they can form the disulfide bridge between chain A to either chain B or chain D (**Fig. 3B**). The neighbouring chain cysteines are positioned so that it is not possible to determine the disulfide bridge orientation. Nevertheless, for modeling purposes, the disulfide bridge was considered to be between chains A and D and between chains B and C. Despite these predicted structures having low PAE and per-chain pTM scores and having different helix conformation compared to the predicted monomer structure, they were selected for further analysis due to their ability to form disulfide bridges (**Supplementary Figure 6**).

### Small-angle scattering studies validate the AlphaFold 3-generated tetramer model

SEC-SAXS analysis of Mam1A_107-213_ (**Fig. 4**; **Table 1**) shows a signature detergent signal as a bump or oscillation at a *q* of 0.18 Å^-1^ in the measured intensities *I*(*q*) (**Fig. 4A**), frequently characteristic of core-shell or micellar structures. Additionally, the Kratky plot suggested a globular shape for the protein (**Fig. 4B**), possibly due to the presence of detergent. The estimated maximal particle distance (*D*_max_) was 90 Å, and the radius of gyration (*R*_g_) was 32.6 Å. The *P*(*r*) curve suggests a primarily globular molecule with an extended region **(Fig. 4C)**. Globular shape is likely partially the result of detergent surrounding the molecule, while the observed extended region could be from the linker region of the expression vector. As the SAXS signal is influenced by the presence of detergent, SANS measurement was conducted on Mam1A_107-213_ (**Fig. 5**; **Table 1**). The detergent signal was contrast-matched (or “masked”) by exchanging the C12E9 into D-C12E9. Three separate samples were subjected to the SANS measurement: i) a sample with non-deuterated C12E9, ii) non-deuterated C12E9 at a CMC concentration, and finally, iii) a sample with deuterated C12E9 **(Fig 5A)**. The signal difference between the three sets gave the first indication that the detergent signal had been matched out successfully. The estimated *R*_g_ and *D*_max_ values for the SANS data were 25.8 Å and 85.0 Å, respectively, which were, as expected, lower than the corresponding values obtained from the SAXS data. Then, we compared the two models against the SAXS data using FoXS [27] and the SANS data using CRYSON[28]. The four-chain arrangement in the predicted model gave the *χ* ^2^ fit of 31.1 for the SAXS data (**Fig. 4D, supplementary Figure 7**). For the SANS data, the fit was much better with a *χ* ^2^ of 2.58 (**Fig. 5C**).

**Fig 4.**
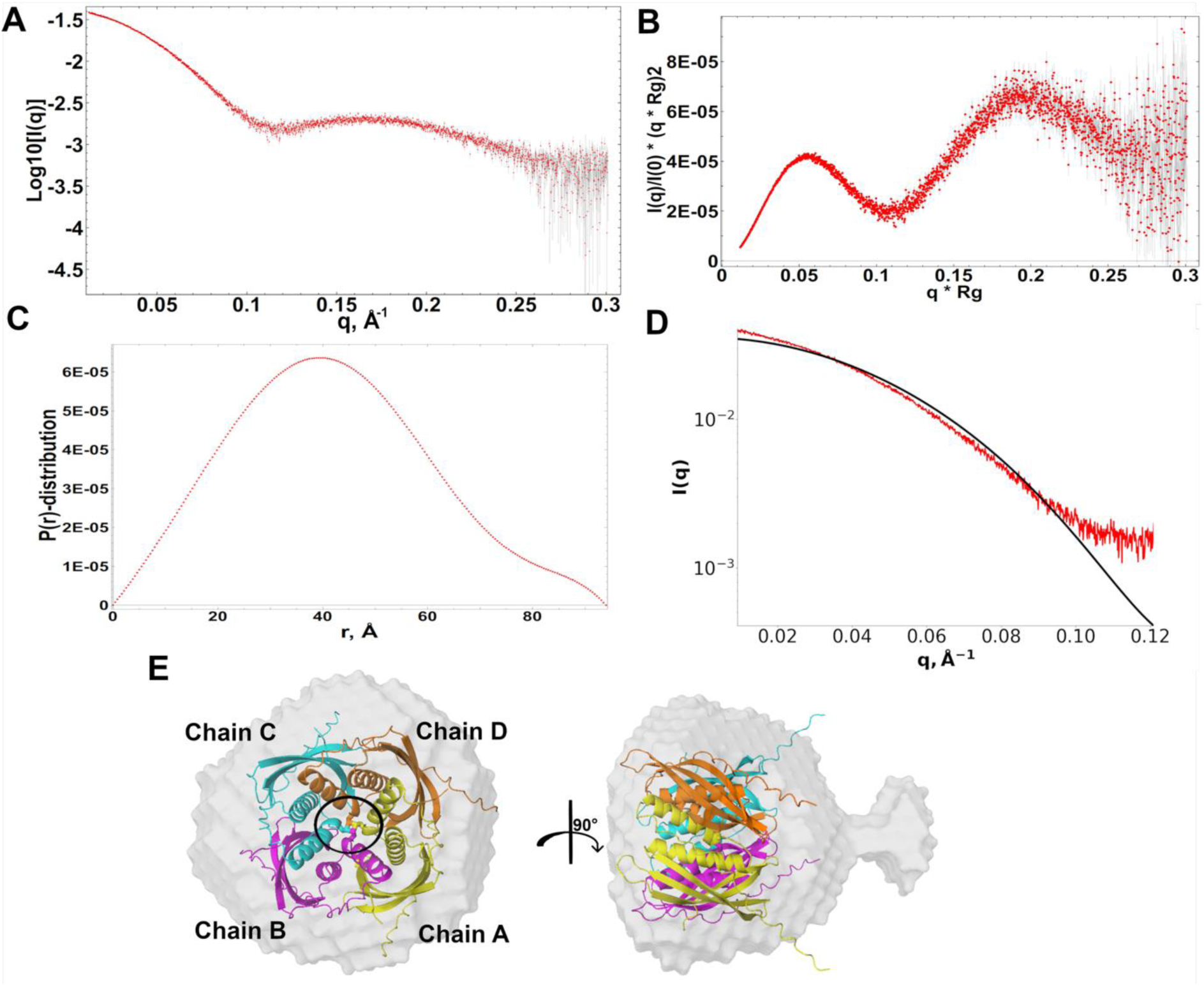
SAXS analysis of Mam1A_107-213_. **(A)** SAXS curve of Mam1A_107-213_. The intensity curve has a characteristic bump/oscillation at *q* of 0.18 Å^-1^ indicating the presence of the detergent micelle with the protein. **(B)** Kratky plot of Mam1A_107-213_ suggesting a folded globular molecule. **(C)** The *P*(*r*) curve of Mam1A_107-213_ SAXS data. **(D)** Fit of the proposed model generated by AlphaFold3 against the Mam1A_107-213_ SAXS data as calculated by FoXS. (E) Superimposition of the tetramer model onto the calculated DAMMIN dummy atom model in a top view and a rotated side view. The disulfide bridge-forming cysteines at the centre of the tetramer are shown as sticks and highlighted in a black circle.

**Fig 5.**
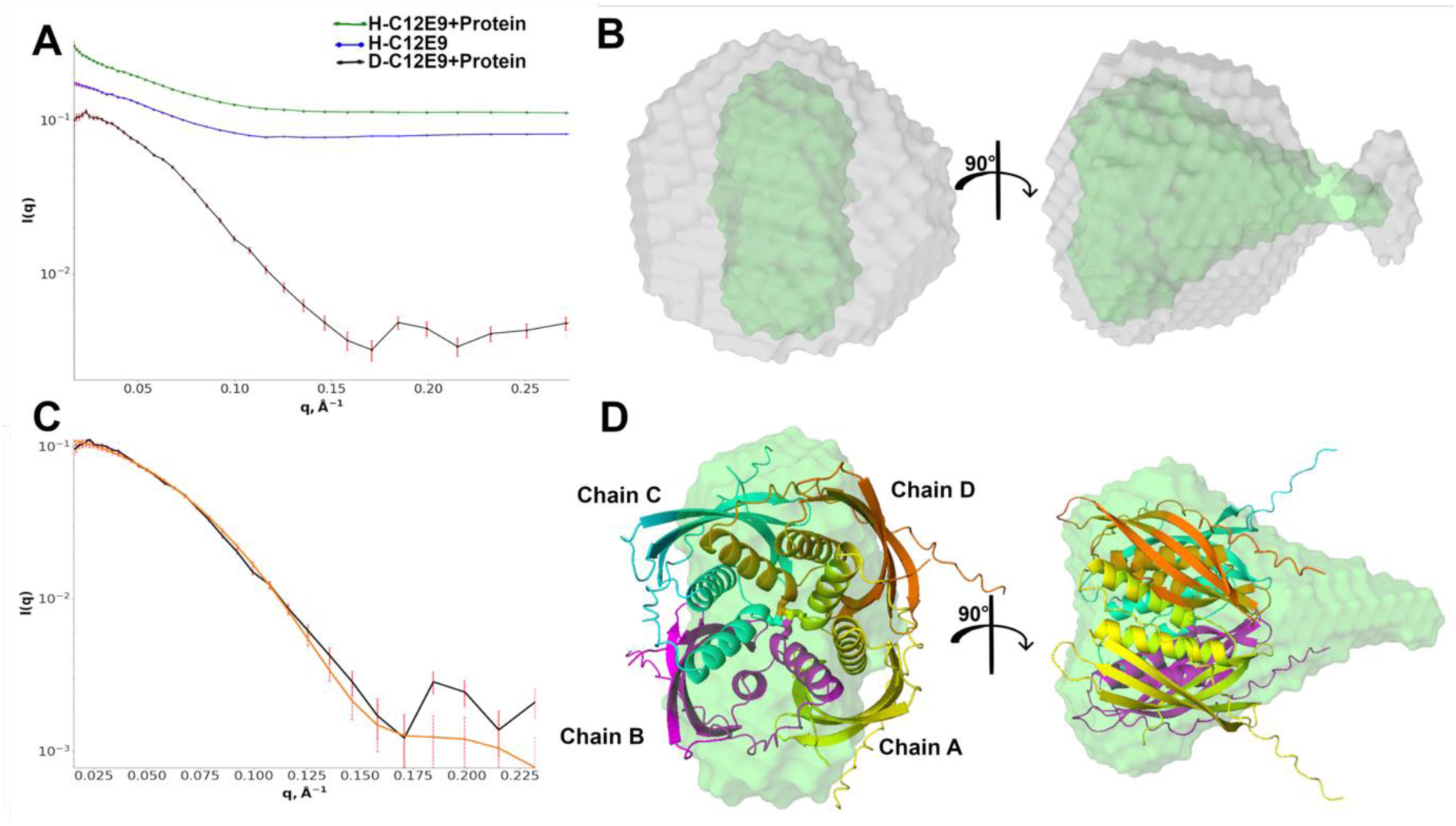
SANS data analysis of Mam1A_107-213_. **(A)** SANS curve of Mam1A_107-213_ in C12E9 (green), C12E9 (blue), and Mam1A_107-213_ in D-C12E9 (black). A comparison of these curves shows that the detergent signal is masked by D-C12E9 resulting in the scattering of only protein molecules. **(B)** Superposition of DAMMIN calculated dummy atoms models from SAXS data (grey) and SANS data (green). **(C)** Fit of the proposed tetramer model against the SANS data as calculated by CRYSON with a *χ*^2^ of 2.58. **(D)** The model superimposed on the calculated SANS dummy atom model in both top and rotated side views.

**Table 1.** SAS data collection parameters and analysis for Mam1A_107-213_.

|  |  |  |
| --- | --- | --- |
| **Protein** | Mam1A<sub>107-213</sub> |  |
| **Source organism** | *Mtb*, H37rv |  |
| **Expression organism** | *E. coli* |  |
| **Experiment** | **SAXS** | **SANS** |
| **Beamline parameters** |  |  |
| Beamline | B21, Diamond Light Source, UK | ZOOM, ISIS Neutron and Muon Source, UK |
| Detector | EigerX 4M(Dectris) | 3He gas tube detector |
| Wavelength, $\text{\AA}$ | 0.9464 | 1.75-16.5 |
| Detector distance, mm | 3732.9 | 4000 |
| $q$-measurement range, $\text{\AA}^{-1}$ | 0.0045 – 0.34 | 0.006 – 0.78 |
| Sample volume, $\mu\text{l}$ | 50 | 300 |
| Sample concentration, $\text{mg ml}^{-1}$ | 8.8 | - |
| **Structural parameters** |  |  |
| $I(0)$ (Guinear), $\text{cm}^{-1}$ | $0.03953 \pm 0.4458 \times 10^{-4}$ | $0.126 \pm 0.9651 \times 10^{-3}$ |
| $R_g$ (Guinear), $\text{\AA}$ | $32.67 \pm 0.4251 \times 10^{-1}$ | $25.8 \pm 0.3431$ |
| $I(0)$ ($P_r$), $\text{cm}^{-1}$ | 0.03953 | 0.126 |
| $R_g$ ($P_r$), $\text{\AA}$ | 32.67 | 25.78 |
| qRg range, Å<sup>-1</sup> | 0.012 – 0.039 | 0.0224 – 0.0458 |
| Limits, qR<sub>gmin</sub> – qR<sub>gmax</sub> | 0.34 – 1.3 | 0.58 – 1.16 |
| q range, Å<sup>-1</sup> | 0.009 – 0.119 | 0.0231 – 0.1708 |
| D<sub>max</sub>, Å | 90 | 85 |
| Porod volume, Å<sup>3</sup> | 177045 | 69729 |
| Mass, kDa (expected) | 74 | 74 |
| Mass, kDa (Porod volume/1.7) | 104 | 41 |
| Mass, kDa (SaxMow2) | 39 | - |
| **Modeling** |  |  |
| Shape modeling software | Primus DAMMIN/DAMAVAR | DAMMIF/DAMFILT |
| q-range for fit (Å<sup>-1</sup>) | 0.009 – 0.119 | 0.0231 – 0.1708 |
| Symmetry assumption | P1 | P1 |
| No. of individual model reconstructions | 20 | 5 |
| Fitting parameters | FoXS | CRYSON |
| χ<sup>2</sup> of the tetramer model | 31.1 | 2.58 |
| **SASDB code** | SASDXF2 | SASDXG2 |

A dummy atom model was generated from the SAXS (**Fig. 4E**) and SANS (**Fig. 5B**) data using the DAMMIN program [29]. The DAMMIN model calculated from the SANS data possessed a different shape than that calculated from the SAXS data (**Fig. 5B**). It should be noted that the SAXS data has a contribution from the detergent and therefore, the generated model based on SAXS data may not be an accurate representation. The SANS model suggests a region of protein forming a tail-like segment pointing away from the main body, suggesting there is a disordered region in the system. A superposition of the fitted tetramer model in CRYSON on SANS-based dummy atom model is shown in **Fig. 5D**.

### Mam1A forms a complex with Mam1C

Following the studies on Mam1A_107-213_, individual expression tests were done for each of the Mam proteins from the *mce1* operon cloned from the *Mtb* genome. Among them, the expressions of Mam1A and Mam1C were observed, both of which belong to the same branch of the phylogenetic tree (**Fig. 1C**). Mam1A and Mam1C were individually purified using His-tag affinity chromatography. Like Mam1A_107-213_, the full-length Mam1A showed the presence of a disulfide bridge, likely involving the same cysteine residue (Cys123). In addition, full-length Mam1C also revealed the presence of a disulfide bridge (**Fig. 6A**). Mam1C has only one cysteine residue in its sequence (Cys46 present in the α1 helix), which is not homologous to the Cys123 of Mam1A, present in the α2 helix. This suggests that Mam1C forms at least a homodimer via a disulfide bridge. A variant of Mam1C was created by removing the first 78 amino acids (Mam1C_79-184_), which are predicted to form the TM helix. Expression and purification of Mam1C_79-184_ showed that this variant is soluble without the need for any detergent and SEC-MALS studies showed that the variant is a mixture of different oligomeric states in solution (**Supplementary Figure 8**). However, the protein started to precipitate at a concentration higher than 1 mg ml^-1^. The variant does not contain the disulfide bridge-forming cysteine, which is probably a contributing factor to the observed instability and heterogeneity. Mam1A and Mam1C were then co-expressed, with His-tag on Mam1A. Purification studies followed by analysis of the bands from SDS-PAGE by mass spectrometry showed that Mam1C co-eluted with Mam1A (**Fig. 6B**). Further analysis of the purified fraction in the presence and absence of BME showed that disulfide bridges were still present in both Mam1A and Mam1C (**Fig. 6B**). There was no indication of a disulfide bridge between Mam1A and Mam1C. These studies suggest that Mam1A and 1C interact to form at least a Mam1A_2_C_2_ heterotetramer (or a higher oligomer).

**Fig 6.**
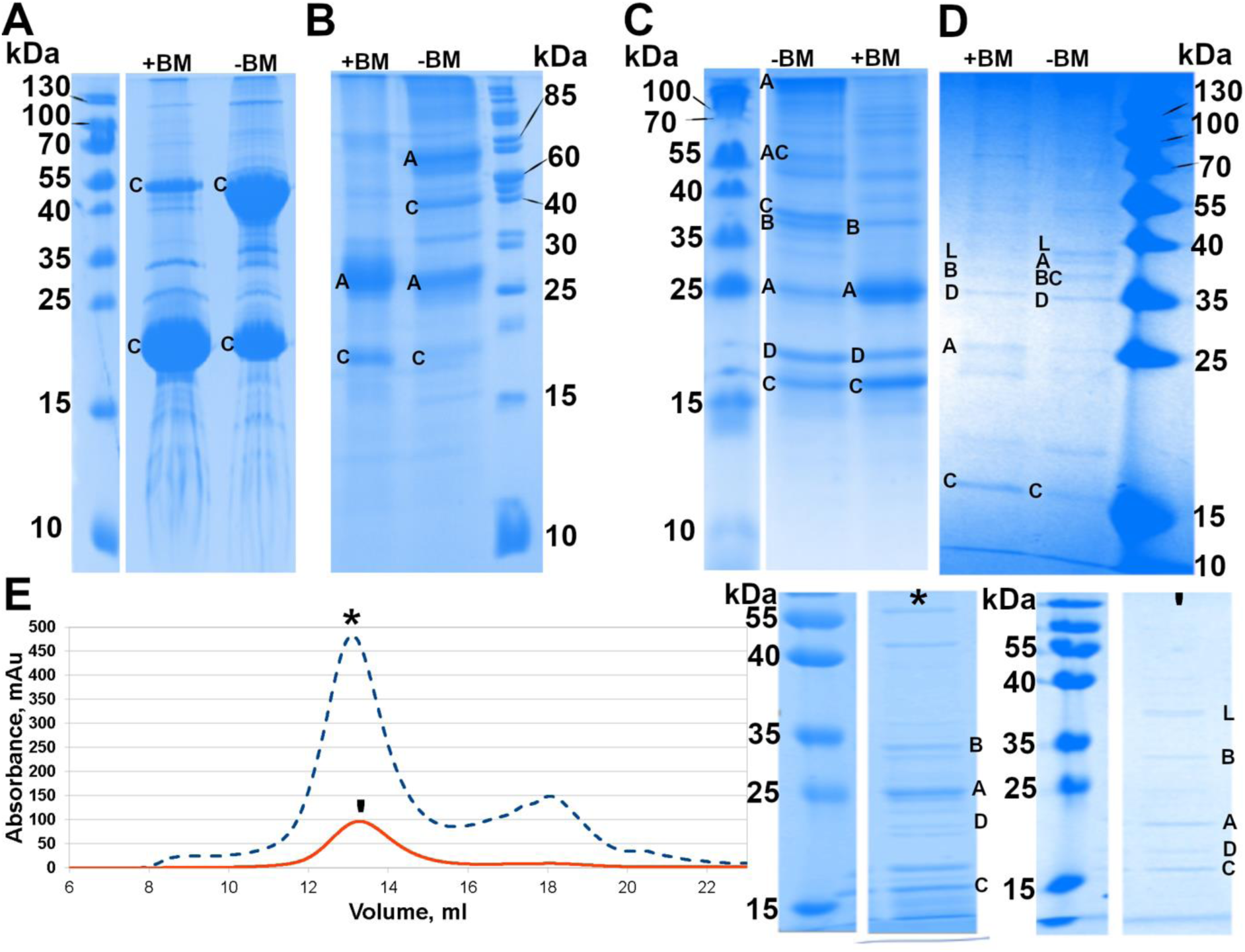
Formation of Mam1AC, Mam1ABCD and Mam1ABCD-LucA Complexes. **(A)** SDS-PAGE analysis of Mam1C in the presence (+BME) and absence (-BME) of BME. The -BME lane has a higher amount of Mam1C at a molecular weight close to 55 kDa showing the presence of an intermolecular disulfide bridge between two Mam1C molecules. **(B)** SDS-PAGE analysis of Mam1A and Mam1C co-expressed and purified showing the formation of the Mam1AC complex. Mam1A contains an N-terminal His-tag. In the -BME, both Mam1A and Mam1C were observed as higher oligomers, suggesting the presence of a disulfide bridge in each of them separately. **(C)** SDS-PAGE analysis of Mam1A-1D co-expressed and purified showing the formation of the Mam1ABCD complex. Mam1A contains an N-terminal His-tag. **(D)** SDS-PAGE analysis of the co-eprxession of LucA with the Mam1ABCD complex. Formation of the Mam1ABCD-LucA complex can be observed. Mam1D contains C-terminal Strep-tag. In SDS-PAGE analysis (C) and (D), both Mam1A and Mam1C were observed as higher oligomers in the absence of BME, suggesting the presence of a disulfide bridge. **(E)** SEC profile of Mam1ABCD (blue dotted line) and Mam1ABCD-LucA (red solid line) and the corresponding SDS-PAGE analysis of the indicated peak fractions. Mam1D contains Strep-tag. The bands in the SDS-PAGE gel corresponding to Mam1A-1D and LucA confirmed by peptide mass fingerprinting analysis are designated as ‘A’, ‘B’, ‘C’, ‘D’ and ‘L’.

### Mam1ABCD forms a stable complex with and without LucA

To test if the other two Mam proteins also interacted with Mam1A and Mam1C, all four Mam1 proteins were co-expressed with Mam1A, with the His-tag on Mam1A. The genes were codon optimized to improve the recombinant expression of these proteins in *E. coli*. Immobilized metal affinity chromatography (IMAC) purification results showed that Mam1B, Mam1C, and Mam1D co-eluted with Mam1A, demonstrating that the four Mam1 proteins interact to form a stable complex. (**Fig. 6C**; **Supplementary Figure 9**). Further, SEC analysis showed two peaks at 13.2 and 16.7 ml in a 24 ml Superose6 column. The SDS-PAGE analysis of the fractions from each of these peaks suggested that both peaks contained all four Mam1 proteins. The intensity of Mam1A was the highest and the intensity of Mam1D was the lowest among the Mam1A-1D in the SDS-PAGE (**Supplementary Figure 9**). As the presence of Mam1D was lower compared to the other Mam1 proteins, a construct was created such that only Mam1D contained a Strep-tag at the C-terminus to have a higher proportion of the Mam1ABCD complex compared to individual proteins. Co-expression and co-purification studies confirmed that Mam1A-1D interact to form a stable protein complex (**Fig. 6E**). With the Strep-tag on Mam1D, the purity and homogeneity of the Mam1ABCD complex improved significantly (**Supplementary Figure 9**).

In a previous study, Nazarova et al.[6] showed that LucA interacts with Mam/Omam proteins using two-hybrid assays. Mam1C is one of the proteins that interacts with LucA. After observing the formation of the Mam1ABCD complex, the interaction of LucA with the Mam1ABCD complex was tested as well. For that, an expression system to co-express all five proteins with His-tag coding sequences on LucA was created. IMAC purification studies showed that LucA co-elutes with other Mam1 proteins, forming a stable Mam1ABCD-LucA complex (**Fig. 6D**). Like the Mam1ABCD complex, the Mam1ABCD-LucA complex showed two peaks in SEC at 12.9 ml and 16.5 ml in a 24 ml Superose6 column (**Supplementary Figure 9**). The expression of LucA containing His-tag produced a more intense band in SDS-PAGE compared to that of the Mam1 proteins (**Supplementary Figure 9**). Therefore, a construct with all five proteins, in which only Mam1D had a Strep-tag, was created. Further Strep-tag affinity purification and analysis of the peak fractions by SDS-PAGE demonstrated the formation of the stable Mam1ABCD-LucA complex (**Supplementary Figure 9**). The purity was higher and the ratio of each of the five proteins was nearly equal using this purification strategy. Mam1A was observed at a slightly lower molecular weight in the SDS-PAGE in the Mam1ABCD-LucA complex compared to Mam1ABCD (**Supplementary Figure 9**). Next, the presence of the disulfide bridges identified in Mam1A and Mam1C during their expression and co-expression was tested in the Mam1ABCD and Mam1ABCD-LucA complexes. Both complexes were analyzed by SDS-PAGE in reducing and non-reducing conditions. The proteins still contained a disulfide bridge and formed homodimers. Both Mam1A and Mam1C were detected as higher oligomers when no reducing agent was used for the analysis of the Mam1ABCD and Mam1ABCD-LucA complexes. These experiments demonstrate that Mam1A-1D can form a stable complex with or without LucA and that LucA interacts with the Mam1ABCD complex. The disulfide bridges were observed in Mam1A and Mam1C proteins as well as in the Mam1ABCD and Mam1ABCD-LucA complexes. Full SDS-page figures of **Fig. 6** can be found in supplementary data (**Supplementary Figure 10 to 15**).

## Discussion

*Mtb* has adapted to switch from using carbohydrates to lipids as the primary energy and carbon source during the intracellular stage of infection [30]. For this, *Mtb* has a large number of lipid-processing enzymes and proteins, including four lipid-importing complexes now known as Mce complexes (Mce1-Mce4). The Mce complexes are shown to import fatty acids, mycolic acids, and cholesterol from the host and its lipid-rich cell wall. The import is not only essential for the survival of *Mtb* in nutrient-depleted environments but also helps *Mtb* to remodel its cell wall in response to its surroundings, playing a critical role in survival, virulence and pathogenicity. Recent studies [6,18,20] have suggested that the Mam/Omam proteins and LucA have a crucial role in the stability and function of the Mce complexes. OmamA is necessary for the function of the Mce4 complex in cholesterol import [18], while LucA similarly stabilizes the Mce1 and Mce4 complexes [6]. The precise mechanism of this stabilization is not yet known. However, the interaction of these proteins with the Mce complexes is likely required. To understand this, we characterized the Mam proteins of the *mce1* operon at a molecular level. The sequence analysis revealed that they have conserved secondary structure features despite having low sequence identity (**Supplementary Table 1**). Further characterization of Mam1A_107-213_ suggested that Mam1A is a tetramer with two disulfide bridges between the pairs of Cys123 when expressed individually. Mutation of this cysteine decreases the protein stability.

Furthermore, we discovered an intermolecular disulfide bridge between Mam1C molecules formed by the sole cysteine Cys46. The disulfide bridge of Mam1C resides in the α1-helix within the TM region of the protein. Deletion of this region (Mam1C_79-184_) produced a soluble variant of the protein. However, the protein was observed to be a mixture of oligomeric states and aggregated during concentration, suggesting the potential importance of a disulfide bridge for its stability. Mam1A and Mam1C interact with each other to form the Mam1AC complex. Furthermore, the four Mam1 proteins and LucA, when co-expressed, form the stable Mam1ABCD and Mam1ABCD-LucA complexes. The disulfide bridges discovered in Mam1A (Cys123) and Mam1C (Cys46) are retained even in the Mam1AC, Mam1ABCD and Mam1ABCD-LucA complexes, highlighting their importance in the overall stability of the complexes. The oligomerization state of Mam1ABCD is thus higher than heterotetrameric. Considering the presence of a disulfide bridge, the minimal unit is likely Mam1A_2_BC_2_D.

As Mam1A_107-213_ was purified in the presence of a detergent, a SAXS measurement alone could not provide all the needed information regarding the shape of the tetramer. Therefore, a SANS measurement performed with Mam1A_107-213_ purified in the presence of deuterated C12E9 to mask the detergent signal via contrast matching, revealing the underlying low-resolution shape information of the tetramer. The dummy atom model produced from the SANS measurement agreed with the geometry of the SAXS measurement yet improved the hypothesis of the protein complex under the detergent micelle. The oligomeric state obtained from the SEC-MALS and the identification of the disulfide bridge played a crucial role in selecting the appropriate model among the predicted AlphaFold3 models. The model in the tetramer had a different conformation of the helices when compared to that in the prediction of individual monomeric Mam1A. Despite these conformational differences and the model showing a low confidence prediction parameter, the fit to the SANS data was excellent, validating and providing insights into the possible assembly of the tetramer. In the proposed model, the disulfide bridges were formed between helices, originating from the neighboring chains.

Previous studies have shown that LucA interacts with Mam1C [6]. It is possible that even in the Mam1ABCD-LucA complex, the interaction of LucA is through Mam1C, which further confirms that the stabilizing effect of LucA to the Mce complexes comes through the interaction with Mam/Omam proteins. Based on our results, we propose that the other Mam/Omam proteins encoded from the same operon can interact to form complexes such as Mam3AB, Mam4AB, OmamAB and OmamCD. It is also possible that the Mam1ABCD complex has a critical role in the stabilization of the Mce1 complex as they are encoded by the same operon. However, LucA, which forms a stable complex with Mam1ABCD, has also been shown to be associated with other Mam/Omam proteins, such as OmamA and Mam4B [6]. This points towards the complex interplay of interaction between different Mam/Omam proteins and their complexes with different Mce complexes. For example, OmamA has been suggested to be essential for the stability of the Mce1 and Mce4 complexes [18]. Our results demonstrate that Mam1A and Mam1C can exist individually and as a part of the Mam1ABCD and Mam1ABCD-LucA complexes. More studies are needed to show if all these forms exist or only as the Mam1ABCD and Mam1ABCD-LucA complexes in the cell. The Mam/Omam proteins are structurally related to VirB8 [18], which is also shown to have multiple interacting partners [31,32]. It is possible that Mam/Omam proteins also behave in a similar manner.

Despite the importance of Mam/Omam proteins and LucA for the functioning of the Mce complexes, these proteins were not present in the recently reported *Mycobacterium smegmatis* Mce1 complex structure [33]. The interaction between Mam/Omam proteins and LucA with the Mce complexes may be dynamic and transient in nature. These interactions are probably necessary at different points in the assembly and functioning of the Mce complexes. Interruption of these interactions can be an effective strategy to destabilize the Mce complexes and pave the way for exploring therapeutic applications for treating latent TB infection. The demonstration and production of the Mam1ABCD and Mam1ABCD-LucA complexes are an important stepping stone for studying their structures and interactions further and provide insights into how these complexes are organized, how they interact with the Mce complexes, and ultimately reveal the mechanism of their action.

## Methods

### Biochemicals

The genomic DNA of *Mtb* H37Rv was purchased from ATCC. Phusion DNA polymerase and the restriction enzymes used for cloning were purchased from Thermo Scientific (Massachusetts, USA) and New England Biolabs. The Ni–NTA chromatography resin for 6xHis-tag purification was obtained from Qiagen (Hilden, Germany). Streptavidin beads for Strep-tag purification were obtained from IBA Lifesciences (Göttingen, Germany).

### Sequence analysis of the genes of interest

Protein sequences were obtained from https://mycobrowser.epfl.ch/. For the sequence analysis and secondary structure predictions, the TMHMM2.0 online server [22], DeepTMHMM1.0 [34], and praline online service tools (Praline) were used. For tertiary structure prediction, AlphaFold2.0 [23] and AlphaFold3.0 [26] were used.

### Cloning *mam1A-1D*, *lucA* and their variants

*mam1A_107-213_* was cloned using the restriction-free cloning method. The gene of interest was amplified from the *Mtb* H37Rv genome using gene-specific primers (forward primer: 5’-<u>AATCTTTATTTTCAGGGCGCCATG</u>GCACCCGATGCTGGGGCG-3’, reverse primer: 5’-<u>GGTGGTGGTGGTGCTCGAG</u>TCACTTCGTCACCTGGTCGAG-3’). *mam1C_79-184_* was cloned similarly using the forward primer: 5’- CTTTATTTTCAGGGCGCCATGGCGTCACCGGATCCGTTTC-3’ and the reverse primer: 5’- <u>GGTGGTGGTGGTGCTCGAG</u>TCAGATCACCGGAAGCAG-3’. The primers from the first PCR reaction created a megaprimer containing the gene and overhangs to the pETM11 recombinant expression vector. The underlined region of the primer is the pETM11-specific sequence. *mam1A_107-213__C123S* was cloned using forward primer 5’- CAGAAGATCATCGAG<u>TCT</u>GGCACCGGTGATTTC-3’ and a reverse primer that was complementary to the forward primer. The underlined region is the mutation site. The mutation was introduced in a single PCR reaction with m*am1A_107-213_* pETM11 as the template by replacing cysteine (TGT) with serine (TCT).

For the co-expression of Mam1A-1D, codon-optimised clones were purchased from GenScript. *mam1A* and *mam1C* were cloned in pET-Duet, while *mam1B* and *mam1D* were cloned in pCOLA-Duet vectors. Altering the protein complex assembly in plasmids was done using the Golden Gate assembly, and *luca* was inserted into the vector with *mam1A* and *mam1C*. Additionally, a pCDF-Duet plasmid containing all the genes of interest, *mam1ABCD* and *luca,* was created using the Golden Gate assembly [35,36].

### Expression and purification of Mam1A_107-213_ and Mam1C_79-184_

The recombinant protein expression was performed in the *E. coli* BL21 (DE3) strain, and the overexpression at 20 °C for 16 h was induced with 0.3 mM to 0.5 mM β-D-1-thiogalactopyranoside (IPTG), once the OD_600_ of 0.6-0.9 was reached. Harvested cells were resuspended in lysis buffer A (500 mM NaCl with 50 mM Tris at pH 8.5) supplemented with 20 µg ml^-1^ of DNase and RNase with an added 0.1 mg ml^-1^ lysozyme and 5 mM MgCl_2_ and lysed by passing the suspension through a cell disruptor at 34.8 kPSi twice. In the case of Mam1A_107-213_, the lysate was incubated with C12E9 (0.1 %) to solubilise the protein. Mam1C_79-184_ was found to be soluble without the use of detergents. The sample was clarified by centrifugation at 32000 *g* for 45 minutes at 4 °C. The supernatant was bound to the Ni-NTA IMAC column and washed with buffer A containing 5 and 50 mM imidazole. The sample was eluted with lysis buffer A with 600 mM imidazole and directly injected into Superdex200 16/600 pg SEC column, which was pre-equilibrated with the appropriate SEC buffer A (500 mM NaCl with 50 mM Tris at pH 8.5). The fractions were analyzed by SDS-PAGE for purity. The pure fractions were pooled, concentrated, and flash-frozen in liquid nitrogen for storage at -70 °C.

### Expression and purification of Mam1AC complex

The codon-optimised constructs were expressed in BL21 (DE3) or BL21 (DE3) RIPL, and the overexpression of proteins was initiated by adding 0.5 mM IPTG to the media once OD_600_ of 0.6-0.9 was reached. Protein overexpression was done at 20 °C for 16 hours. The Mam1AC complex purification followed the same protocol as Mam1A purification, with 5 mM FosCholin-12 (FC12) supplementation to the buffer. Mam1AC purification required detergent in the following steps of IMAC and SEC, for which the buffers were supplemented with 0.05% C12E9.

### Expression and purification of Mam1ABCD and Mam1ABCD-LucA complexes

The expression of the Mam1ABCD and Mam1ABCD-LucA complexes was done in the Rosetta (DE3) pLysS strain. The expression was initiated with 1 % primary culture in the terrific broth-auto induction medium (TB-AIM) and grown at 37 °C until OD_600_ of 0.25-0.3 was reached, after which the temperature of the media was brought down to 22 °C and expression was allowed to proceed for 20-22 hours. The cells were harvested by centrifugation and resuspended in lysis buffer B (500 mM NaCl and 10 % glycerol with 50 mM Tris at pH 8.0) supplemented with 20 µg ml^-1^ of DNase and RNase, 0.1 mg ml^-1^ lysozyme, 10 mM MgSO_4_, 25 mM DDM, one tablet Protease COCKTAIL (Catalog #S8830 Sigma Aldrich), and 25 mM ATP. The His-tag purification was continued by supplementing lysis buffer B with 5 mM imidazole. The cells were lysed by sonication, and cell debris was removed by centrifugation at 28800 *g* for 40 minutes at 4 °C. IMAC purification was initiated by incubating the supernatant with Ni-NTA beads equilibrated in lysis buffer B supplemented with 1 mM DDM and 5 mM imidazole for one hour. The sample was eluted by adding elution buffer (500 mM NaCl, 10 % glycerol, 1 mM DDM and 500 mM imidazole with 50 mM Tris at pH 8.0) to the beads. In the case of Strep-tag purification, the cells were resuspended in lysis buffer B with the same supplementation except for the imidazole, and they were similarly lysed and clarified as in Ni-NTA purification. The complex was bound to Strep-beads and eluted in lysis buffer B supplemented with 70 mM biotin. In the case of both Ni-NTA and Strep-tag purification, the eluted flow-through was concentrated by centrifugation to a volume of around 1 ml using 100 kDa cutoff centricons. The concentrated sample was injected into the Superose6 10/300 GL SEC column, equilibrated in SEC buffer B (500 mM NaCl, 5 % glycerol and 1 mM DDM with 50 mM Tris at pH 8.0). The peak fractions were analyzed by SDS-PAGE, and the identity of the bands was confirmed by peptide mass fingerprinting analysis using MALDI-TOF MS.

### Unfolding of Mam1A_107-213_ by NanoDSF

The instance of aggregation as well as *T_m_* was calculated at a concentration of 0.1 mg ml^-1^, 0.3 mg ml^-1^, and 0.6 mg ml^-1^ from 20 °C to 90 °C with 1 °C per minute temperature increase, using standard NanoTemper Prometheus capillaries (PR-C002). NanoDSF data were analyzed using the MoltenProt online service program [25,37].

### Polydispersity analysis of the purified Mam1A_107-213_ by DLS

The polydispersity of the sample at concentrations 0.2 mg ml ^-1^, 0.4 mg ml ^-1^ and 0.6 mg ml^-1^ in triplicates was measured by DynaPro Platereader-II and analyzed using Dynamics version 10.1.0.

### Mam1A_107-213_ SEC-MALS analysis

50 µl of purified protein at 5 mg ml^-1^ was injected into a Superdex 200 10/300 GL column, connected to a Shimadzu purification system, a Shodex refractive index detector, and a Wyatt mini-DAWN Treos MALS detector. The results were analyzed by Astra 7.0 software. UV and RI measurements were used to calculate the molecular weight of the protein complex.

### Mam1A_107-213_ SEC-SAXS sample preparation, data collection and processing

SAXS measurements were carried out at the Diamond Light Source. 50 to 100 µl of the sample at concentrations ranging from 1 to 8 mg ml^-1^ were injected into a UHPLC coupled system (SEC-SAXS). An inline SEC-SAXS sample was injected into a Superdex200 3.2/300 GL column pre-equilibrated in SEC buffer A. Measurement of the sample was done with PILATUS 2M two-dimensional with a sample to detector distance of 3732.9 m at a wavelength of 0.9464 Å at the beamline B21 in Diamond Light Source UK.

SAXS data analysis and buffer subtraction were done using the Scåtter III and IV programs[38,39]. With the Scåtter *R*_g_, *D*_max_ and forward scattering *I*(0) were calculated. The produced output file was used for online ATSAS services to model the DAMMIN dummy atom model [29,40]. The predicted AlphaFold2.0 and AlphaFold3.0 models [23,26] were then fitted against SAXS data using the FoXS online server [27].

### Preparation of deuterated C-12E9 (D-C12E9)

Tetraethylene glycol (25 g) was dried under vacuum at 60 °C for 24 hours. *t*-BuOK (1.11 g, 10% excess) was then added, and the mixture was stirred under vacuum at 60 °C for 2 hours. Subsequently, 1-bromododecane (3 g) was introduced, and the reaction mixture was heated to 110 °C under vacuum for 2–3 hours. After completion, a few drops of HCl were added to neutralize the reaction, followed by extraction with ether. The crude product was concentrated via rotary evaporation, yielding 4.94 g before purification by flash column chromatography.

The chain-deuterated species were synthesized using the same procedure as the protonated counterpart, replacing 1-bromododecane with 1-bromododecane-d₂₅. Deuteration was achieved from dodecanoic acid, where 25 g of protonated dodecanoic acid was mixed with 10 % Pt/C (10 wt %) in 300 mL of D₂O and subjected to a Parr reactor at 180 °C for three days. After two H/D exchange cycles, the deuteration level was confirmed by nuclear magnetic resonance (NMR) (**Supplementary Figure 16**). The resulting deuterated dodecanoic acid was then reduced to the corresponding alcohol using LiAlD₄. Bromination was performed by reacting the deuterated alcohol with 48 % HBr in the presence of TBAB as a phase transfer reagent. The product was extracted with diethyl ether, concentrated under vacuum, and purified via column chromatography.

### Mam1A_107-213_ SANS sample preparation, data collection and analysis

200 µl of protein sample at the concentration of 10 mg ml^-1^ were mixed with 0.2 % deuterated C12E9 (D-C12E9) suspended in 50 mM Tris with 500 mM NaCl at pH 8.5 in a 1:1 ratio. D-C12E9-protein mixture was kept on ice for an hour before injecting the sample into the Superdex 200 10/300 GL column. The column was pre-equilibrated in SEC buffer with twice the critical micelle concentration (CMC) of C12E9 (CMC = 0.003 %). For the experiment, it was assumed that the CMC of D-C12E9 was the same as that for H-C12E9.

SANS Data were collected on the Zoom time-of-flight SANS instrument (ISIS Neutron and Muon Source, Harwell, UK) using a wavelength range 1.75-16.5Å. The source to sample and sample to detector distance was 4 m and the beam size was 12 mm. From the highest point of the peak, 3 fractions were selected. Each of these fractions was loaded into a Starna Scientific quartz round cuvette (match code 6, path length 1, type 32/Q/1) and SANS data were collected from each fraction with an exposure time of 1 h in 5 cycles (5 h total exposure time for each fraction) at 6 °C in a temperature-controlled sample rack. The detector images were reduced to 1-D using Mantid software, including normalization, detector correction, absolute scaling and buffer subtraction. SANS data were processed using the same method outlined above for the SAXS data. The predicted AlphaFold3 structures were fitted against the SANS data using the CRYSON from online ATSAS^43^.

## Acknowledgement

The work is supported by funding from the Tampere Tuberculosis Foundation, Research Council of Finland (332 967 & 371 075), the University of Oulu Graduate School, and the Jane and Aatos Erkko Foundation. We acknowledge the support from Diamond Light Source, UK, for SAXS data collection and ISIS Neutron and Muon Source, UK, for SANS data collection. We also acknowledge the support from the Biocenter Oulu structural biology, proteomics and protein analysis, and DNA sequencing core facilities.

## Data availability

The relevant data associated with the published study are present in the paper or attached as Supplementary data. The SAXS and SANS data are available with the codes SASDXF2 and SASDXG2, respectively, from sasdb.org.

## Author contributions

R.V. with M.J.H. planned the study. M.J.H. executed the cloning, purification, and characterization of Mam1A, Mam1C, Mam1AC and their variants. P.P. and N.T.H. created the Mam1ABCD and Mam1ABCD-LucA expression constructs. P.P. and M.J.H. executed the purification and identification of Mam1ABCD and Mam1ABCD-LucA-related complex profiling. K.M. prepared the C12E9 and D-C12E9 for the SANS experiment. J.J.D. assessed the SANS data and modeled the respective dummy atom model from the dataset. R.V., M.J.H. and N.T.H. wrote the manuscript with contributions from other authors. R.V. acquired the funding for the work.

