## Supplementary Information for "Insight into the structure and interactions of the *M. tuberculosis* Mce-associated membrane proteins Mam1A-1D"

### Supplementary Figures

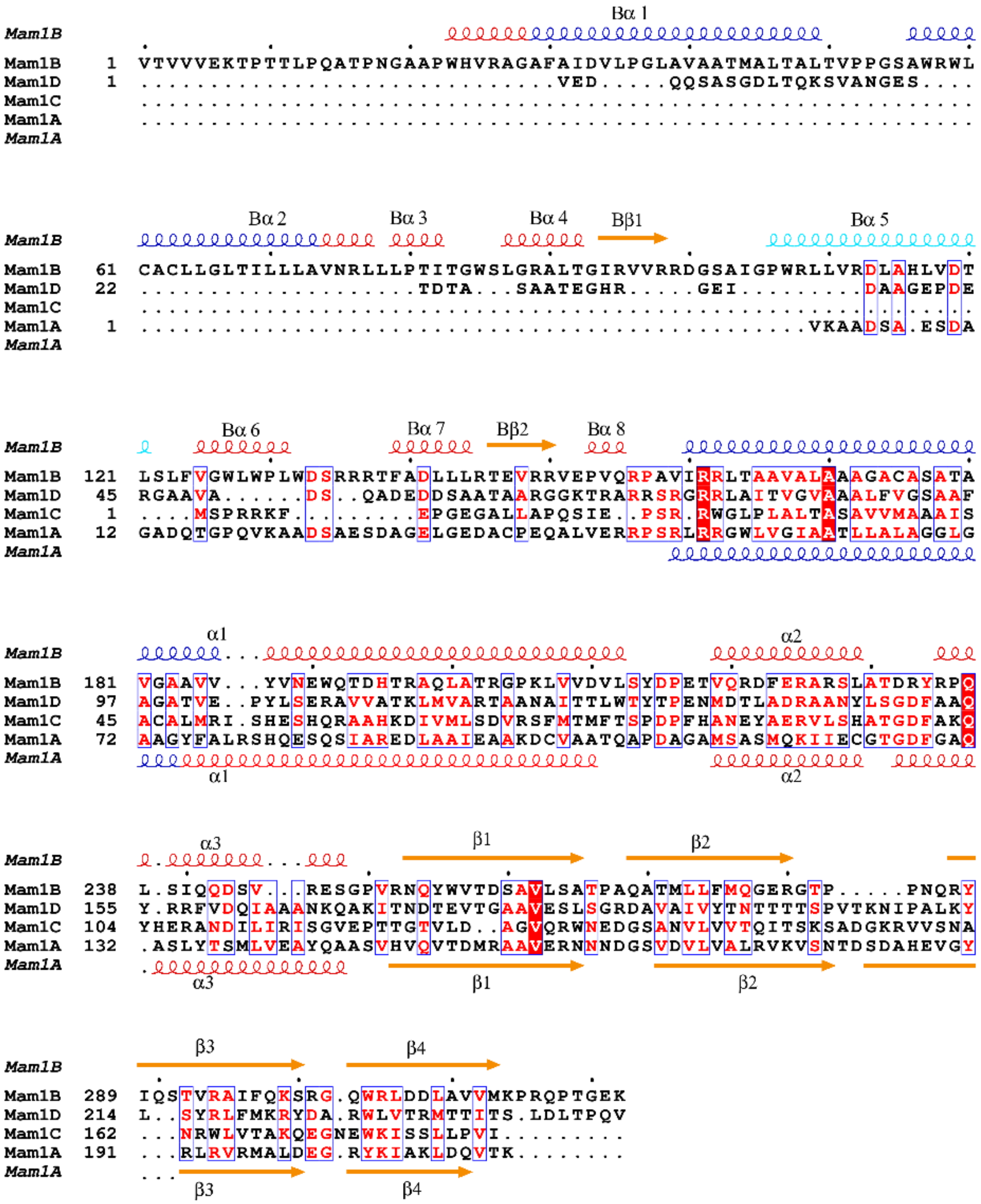


**Supplementary Figure 1. Sequence alignment of Mam1 proteins.** Sequence alignment of Mam1 proteins shows that there are only four amino acids conserved among the proteins. However, the structural elements are similar, excluding the variable region. Secondary structure elements from AlphaFold2.0 predictions are shown for Mam1A (bottom) and Mam1B (top).

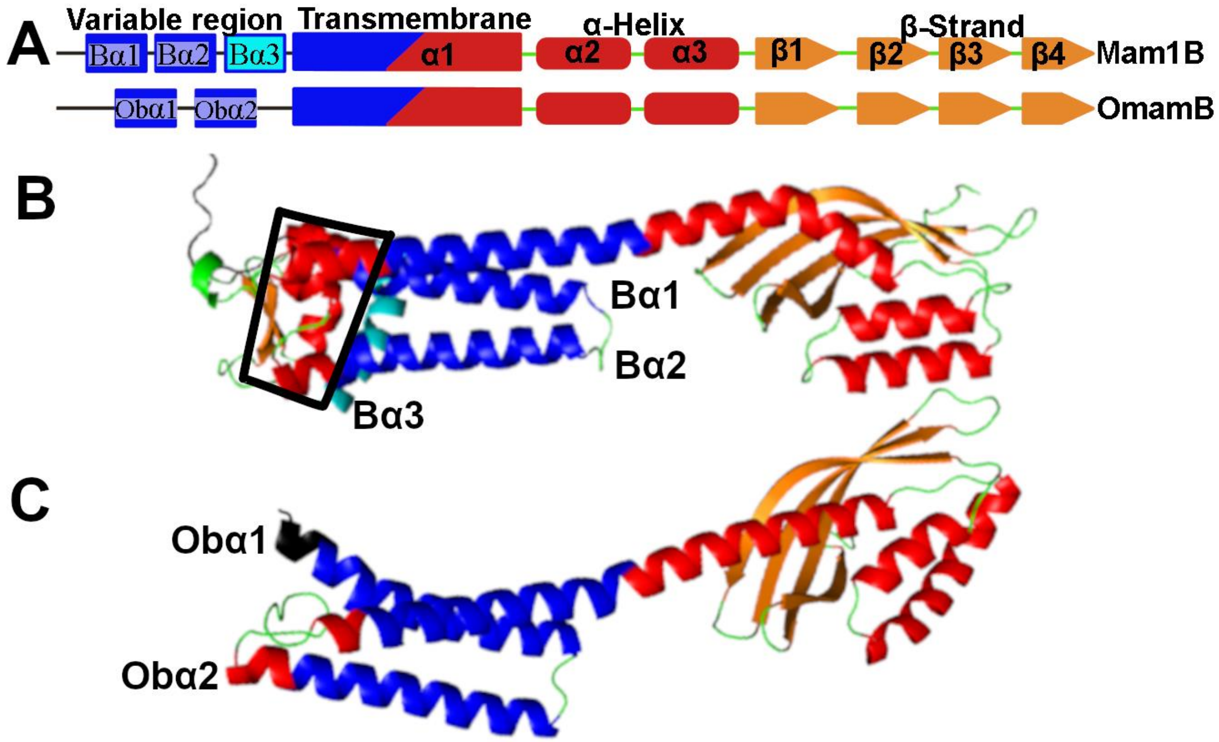


**Supplementary Figure 2. Secondary structure elements and AlphaFold prediction of Mam1B and OmamB**. **(A)** A schematic illustration of the general conserved secondary structure organization of Mam1B and OmamB. The four regions are named i) variable region, ii) transmembrane, iii) α-helix and iv) β-strand. Unlike the rest of the Mam and Omam proteins, Mam1B and OmamB have three transmembrane helices. **(B)** Predicted structure of Mam1B is shown to have an RDD domain (black box) that resides in the N-terminal region of the protein. This region is also the variable region in Mam1B. Mam1B additionally has an α-helix, that might be buried in the membrane, annotated as Bα3 and colored in cyan. **(C)** Predicted structure of OmamB. The variable region in OmamB is relatively short (helix region colored in black), but the protein does contain three TM-helices, like Mam1B.

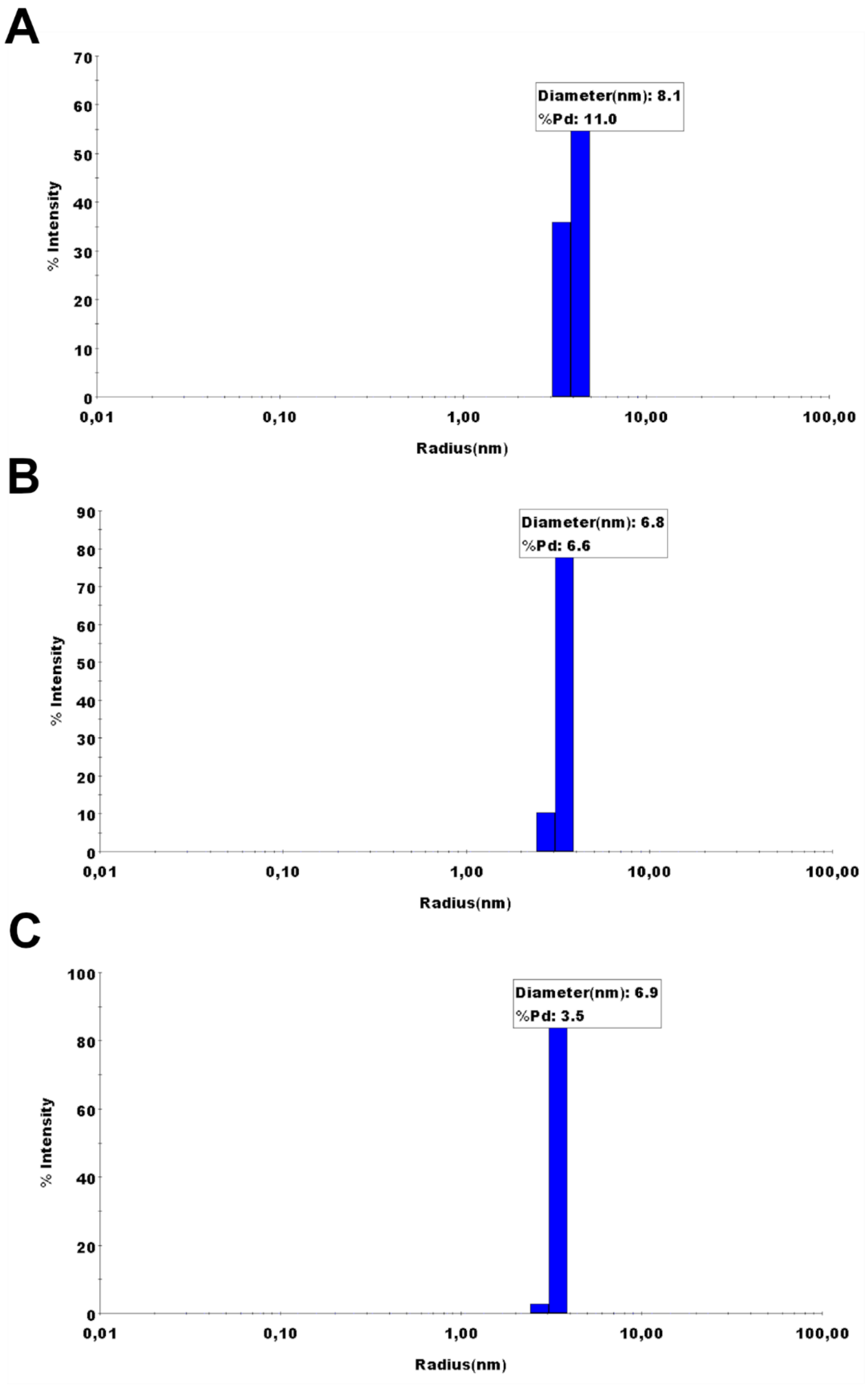


**Supplementary Figure 3. Polydispersity and particle diameter estimation of Mam1A$_{107-213}$ using DLS measurement.** The polydispersity was measured in three concentrations. **(A)** 0.2 mg ml$^{-1}$, **(B)** 0.4 mgml$^{-1}$ **(C)** 0.6 mgml$^{-1}$ in 0.05% C12E9

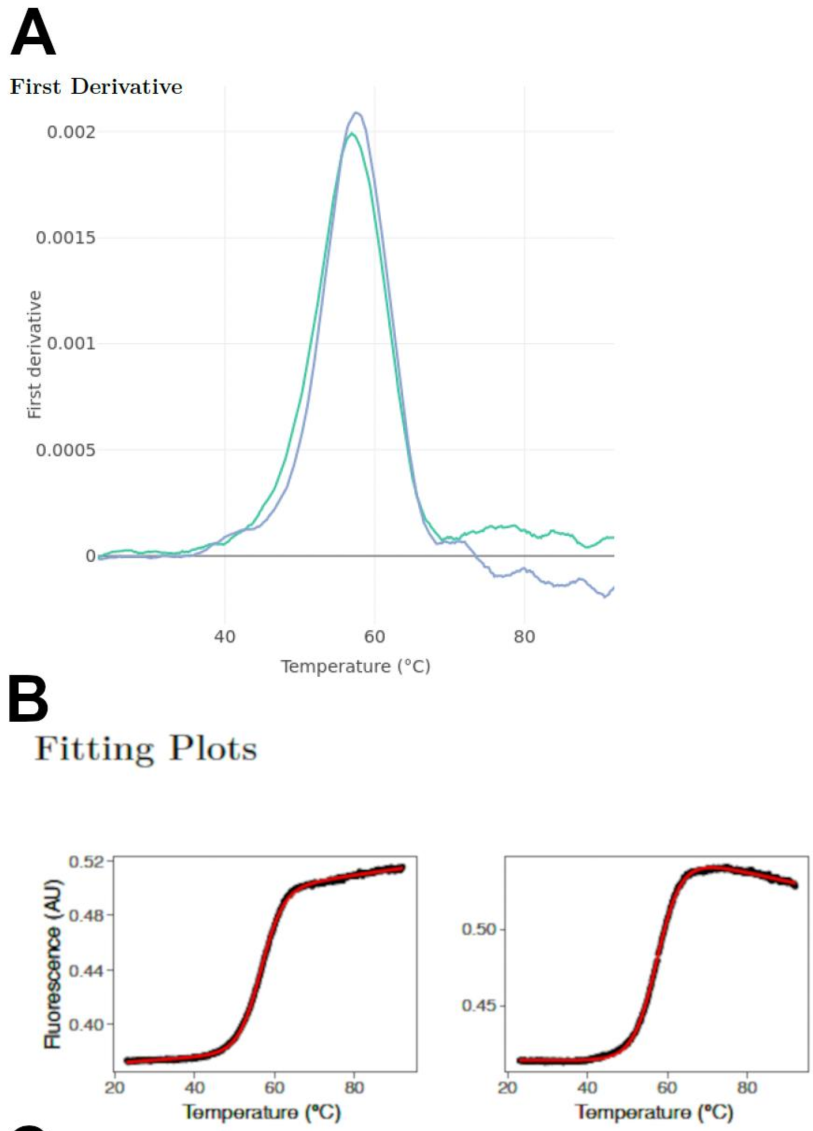


**C**

Fitted Parameters

| kN | bN | kU | bU | dHm | Tm |
|---|---|---|---|---|---|
| 0.000231 | 0.304 | 0.000417 | 0.362 | 75.9 | 56.5 |
| -4.29e-06 | 0.416 | -0.000569 | 0.738 | 71.5 | 57.6 |

**Supplementary Figure 4**. **Moltenprot analysis of Mam1A$_{107\text{-}213}$from two separate purifications. (A)** First derivative plot of the measured melting curve. Two separately purified samples were measured for consistency. **(B)** Fitting(red) of the measured melting curve(black). **(C)** Fitting parameters and calculated values of the melting curve produced by the MoltenProt webservice.

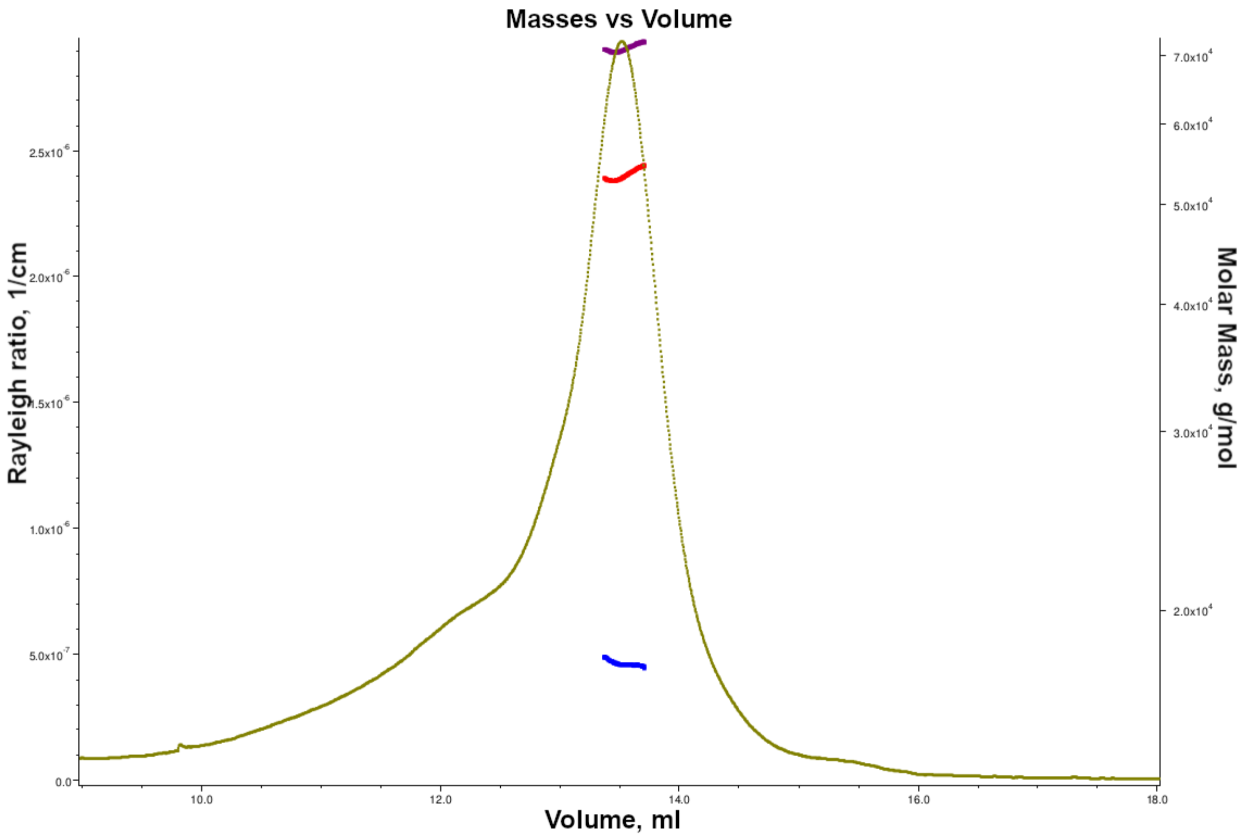


**Supplementary Figure 5. SEC-MALS protein-detergent analysis of the Mam1A$_{107\text{-}213}$ sample with C12E9.** The SEC-MALS measurement of Mam1A$_{107\text{-}213}$ shows the protein-detergent complex formation. Plotted curve is the measured light scattering (LS) signal of the sample. Purple line (topmost) indicates the total mass of the complex, red line (middle) indicates the calculated protein mass, and blue line (bottommost) indicates the calculated detergent mass. For the protein conjugate analysis, the dn/dc value of C12E9 was taken from anatrace (0.109 ml/mg).

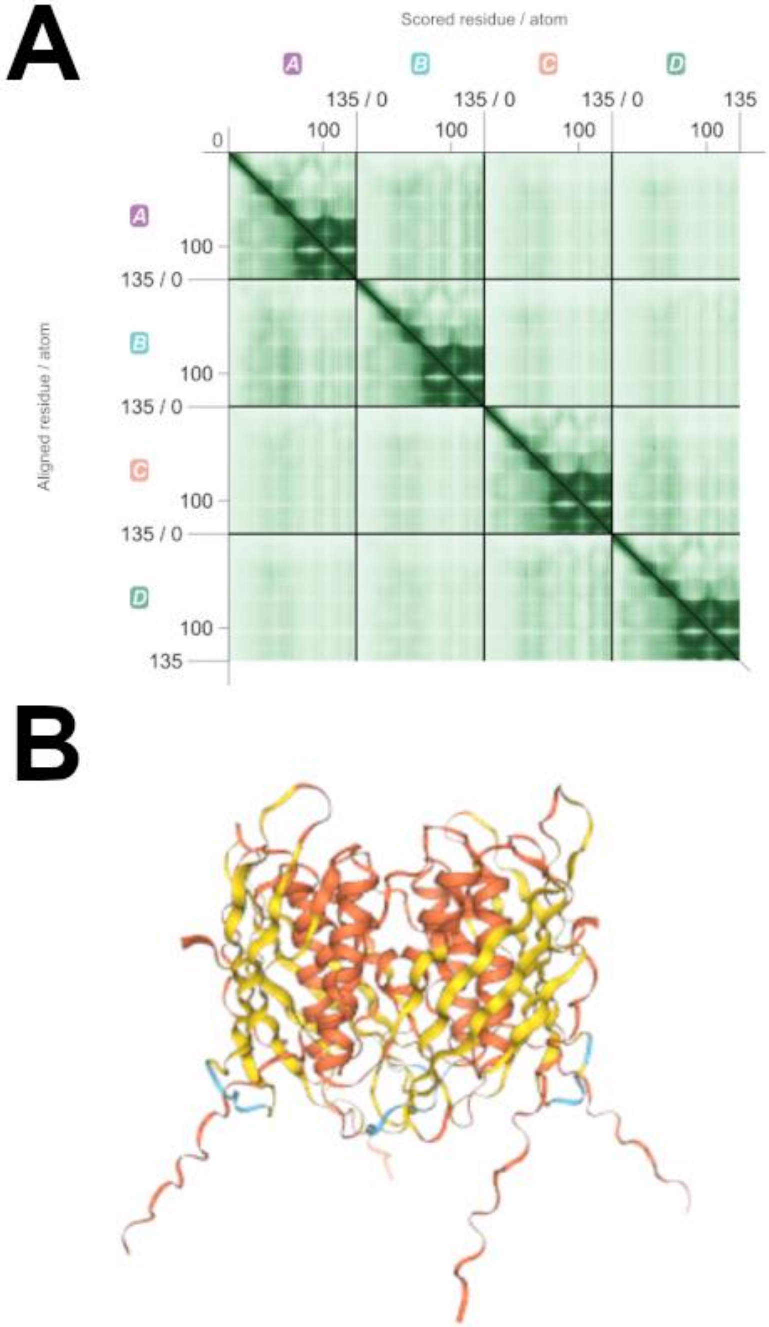


**Supplementary Figure 6**. **Prediction quality parameters of Mam1A$_{107\text{-}213}$ tetramer. (A)** AlphaFold3-predicted aligned error map of Mam1AΔ106 tetramer. **(B)** AlphaFold pLDDT color-coded scoring by residue. Image generated with PAE Viewer from a JSON file produced by AlphaFold3.

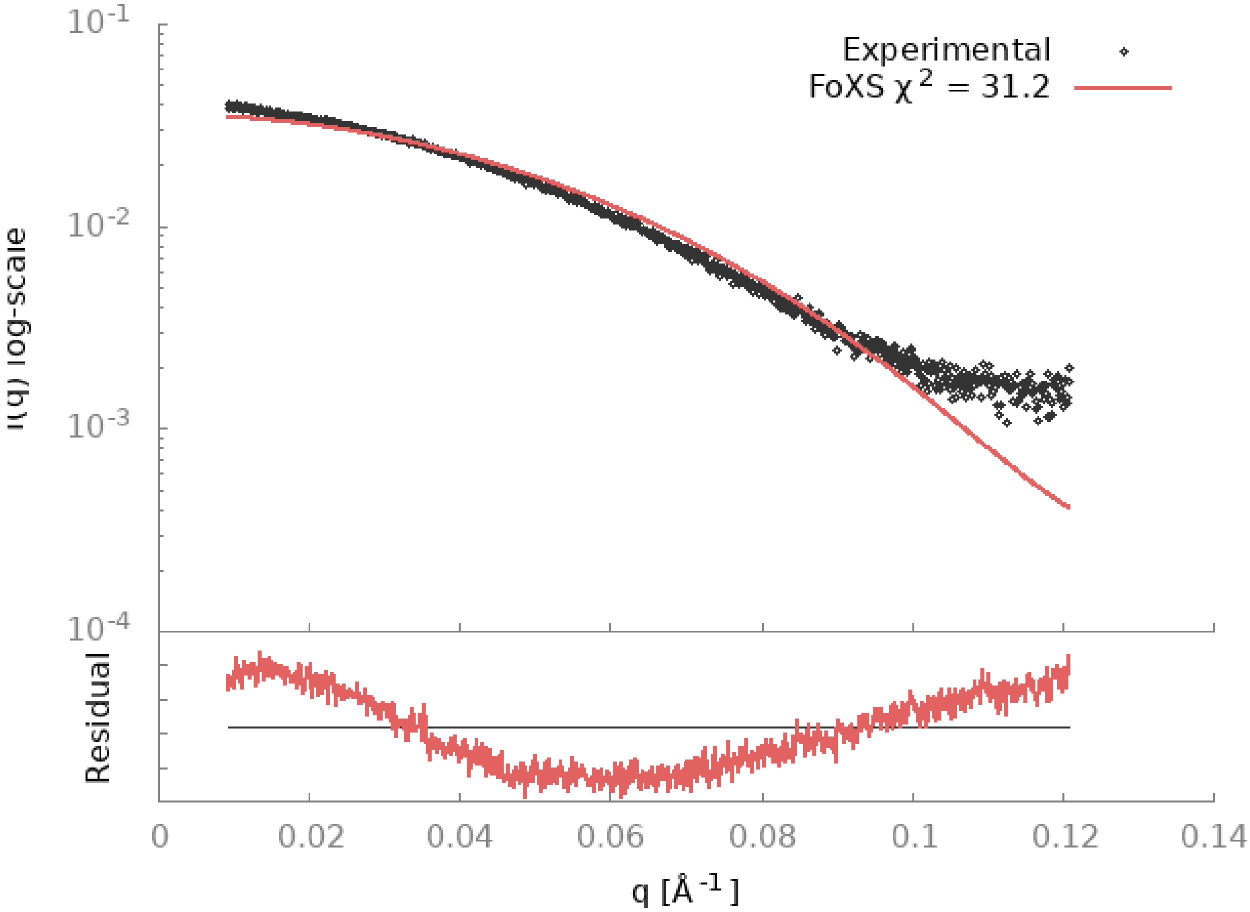


**Supplementary Figure 7. Tetramer model fit of Mam1A$_{107\text{-}213}$ to the SAXS data.** Tetramer model $\chi^2$ score and the fit of the model to the SAXS data. Residual fit of the model to the SAXS data has a relatively poor fit, likely caused by the fact that the detergent is not modeled. Image produced by the FoXS webservice.

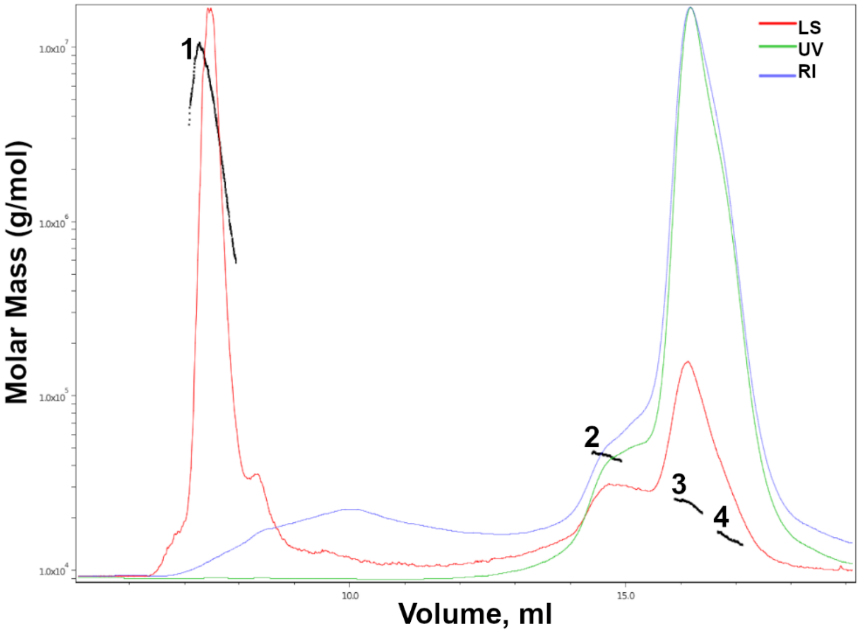


**Supplementary Figure 8. SEC-MALS analysis of the Mam1C$_{79-184}$ variant.** The SEC-MALS analysis of the soluble Mam1C$_{79-184}$ variant indicates that the variant is mostly in solution as a monomer but also forms a higher oligomer. Masses calculated for the indicated fractions were 1.6 MDa(1), 45 kDa (2), 23 kDa (3), and 15 kDa (4). Theoretical size for the variant is 15 kDa. The variant was noted to aggregate even at lower concentrations. The sample was injected into the SEC-MALS column at a concentration of 1 mg ml$^{-1}$.

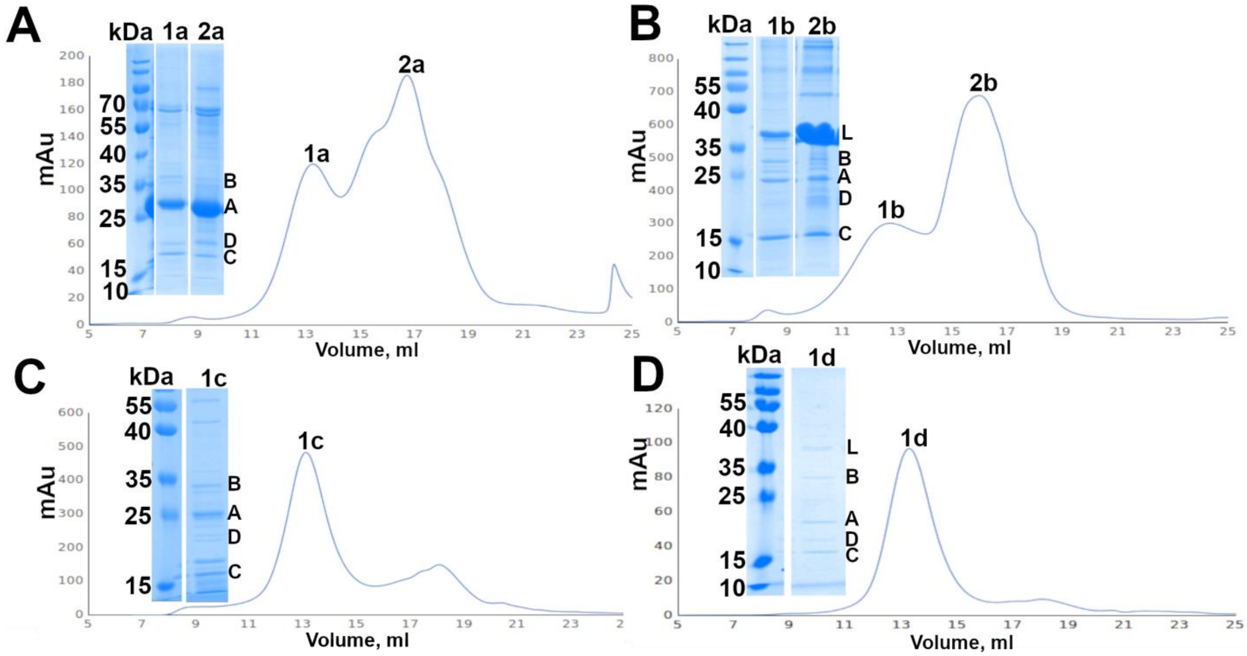


**Supplementary Figure 9. Mam1ABCD purification profiles with and without LucA with different purification tags.** The Mam1ABCD complex was purified without (**A** and **C** profiles) and with LucA (**B** and **D** profiles). **(A)** Co-purifcation of Mam1ABCD with a purification tag in Mam1A and **(B)** Luca which produced two peaks on the SEC purification. Peak #2 has substantially more of the tagged protein, while both peaks harbor the full complex. **(C)** Co-purification of Mam1ABCD complex with strep-tag on Mam1D, showing a more uniform distribution of Mam1A-1D present. **(D)** Mam1ACBD-LucA co-purification, with purification tag in Mam1D, resulting in a single major peak in SEC.

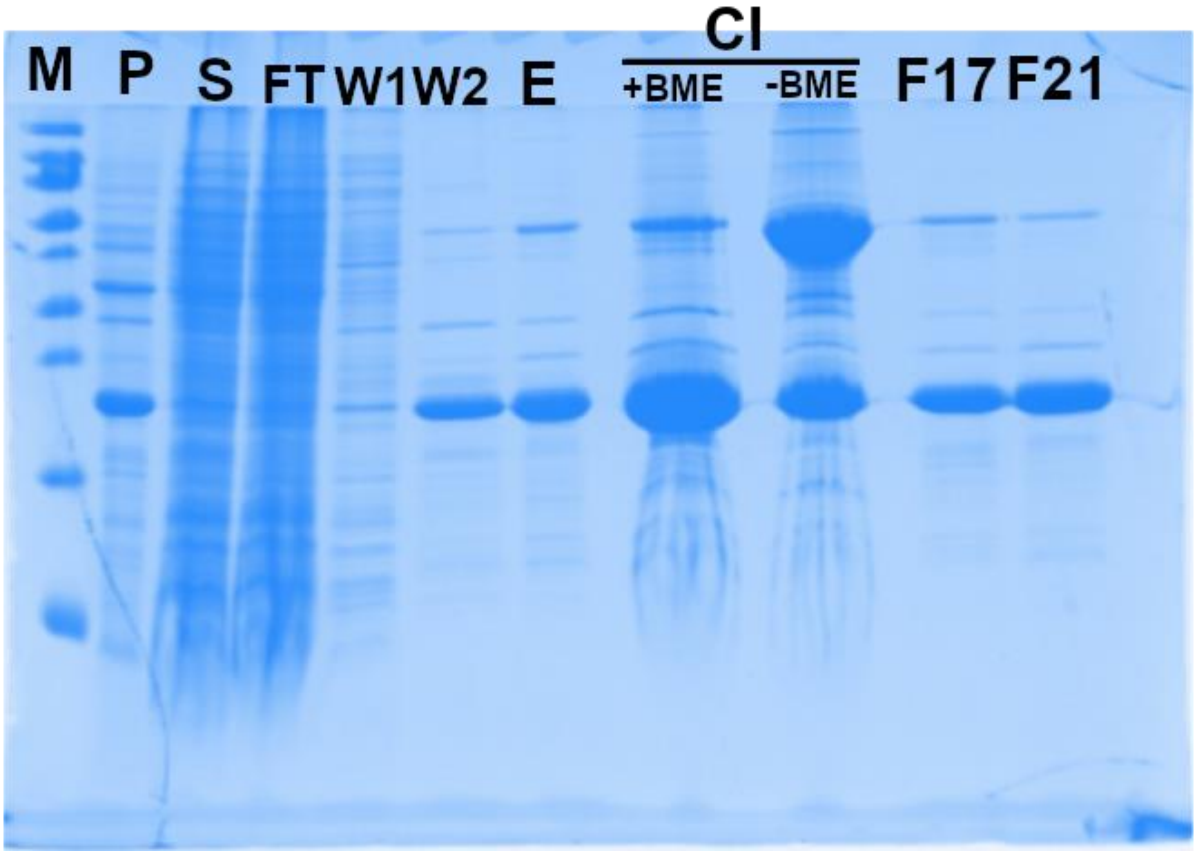


**Supplementary Figure 10**. Full SDS-PAGE figure of supplementary **Fig. 6A**. Mam1C purification. Analysis with and without BME. Abbreviations: M = Standard protein ladder, P = pellet fraction from centrifugation after cell disruption, S = supernatant from centrifugation after cell disruption, FT = IMAC flow through, W1 = 1st wash step flow through of the IMAC, W2 = 2nd wash step slow through of IMAC, E = Elute sample of the IMAC, CI = Concentrated sample injected to SEC, BME = β-mercaptoethanol, F12 and F21 are two fractions from the observed peak.

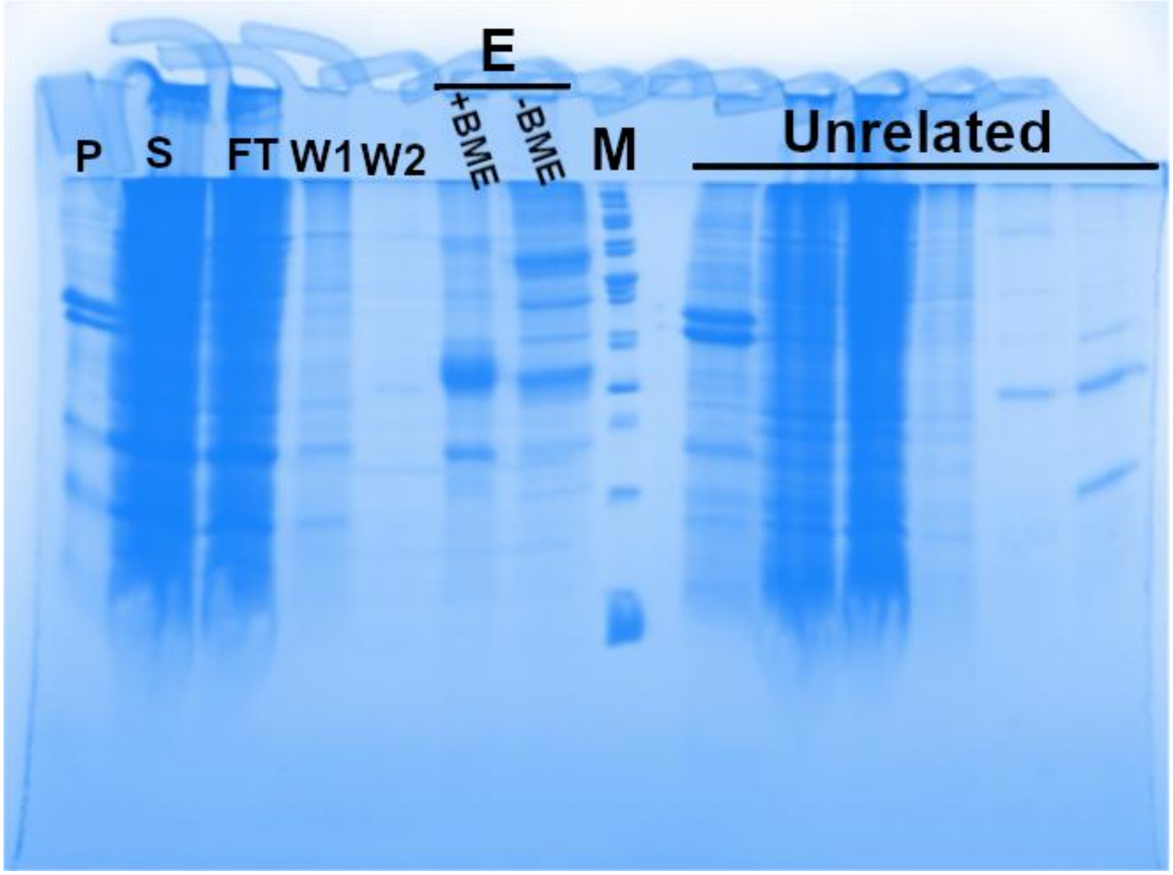


**Supplementary Figure 11**. Full SDS-PAGE figure of supplementary **Fig. 6B**. SDS-PAGE analysis of affinity purification of Mam1AC with His-tag at the N-terminus of Mam1A. The eluted protein was run with and without β-mercaptoethanol (BME). Abbreviation: P = pellet fraction from centrifugation after cell disruption, S = supernatant from centrifugation after cell disruption, FT = flow through of IMAC binding step, W1 = 1st wash step flow through, W2 = 2nd wash step flow through, E = elution fraction from affinity step, with and without BME, M = standard protein ladder. The right side of the gel was used for unrelated protein purification, not discussed in this paper.

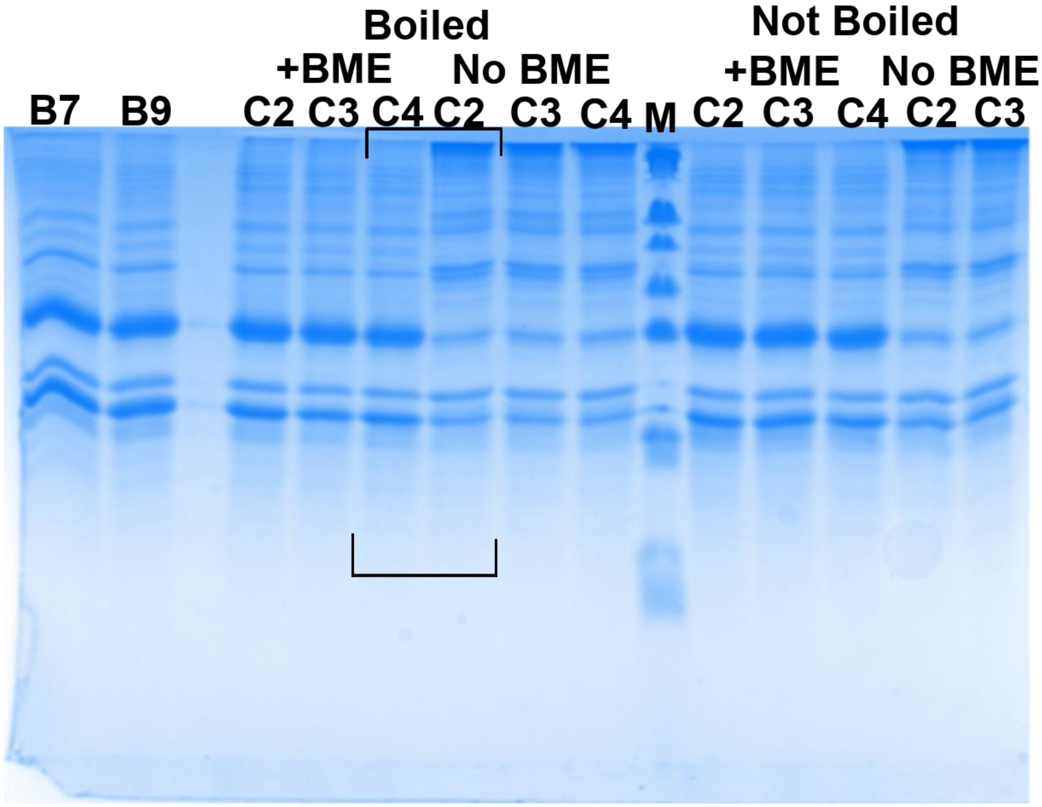


**Supplementary Figure 12**. Full SDS-PAGE of **Fig. 6C**. Mam1ABCD Äkta-purification fractions, triplicates, with and without BME and with and without boiling the sample at 100 °C for 10 minutes. A His tag was placed in Mam1A. Brackets indicate the area shown in **Fig. 6C**. Fraction names corresponding to the volume and peak: Aggregate peak: B7=9ml, B9=10ml. Main peak: C2=13 ml, C3=13.5 ml, C4=14 ml.

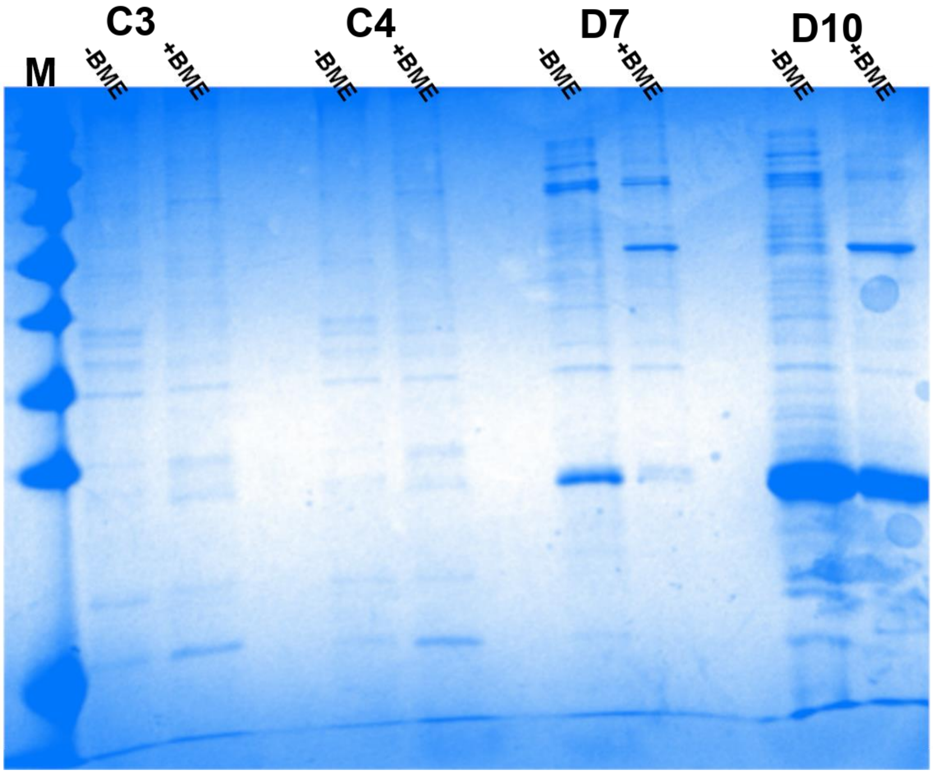


**Supplementary Figure 13.** Full SDS-page of **Fig. 6D**. Mam1ABCD+LucA Superose 6 gel filtration duplicate fractions with and without β-mercaptoethanol (BME). A Strep tag was placed in LucA. Fraction C3 was shown in **Fig. 6D**. Fraction names corresponding to the volume and peak: C3=12.1 ml, C4=12.4 ml (main peak 2), D7= 17.2 ml D10=17.8 ml (late peak 3)

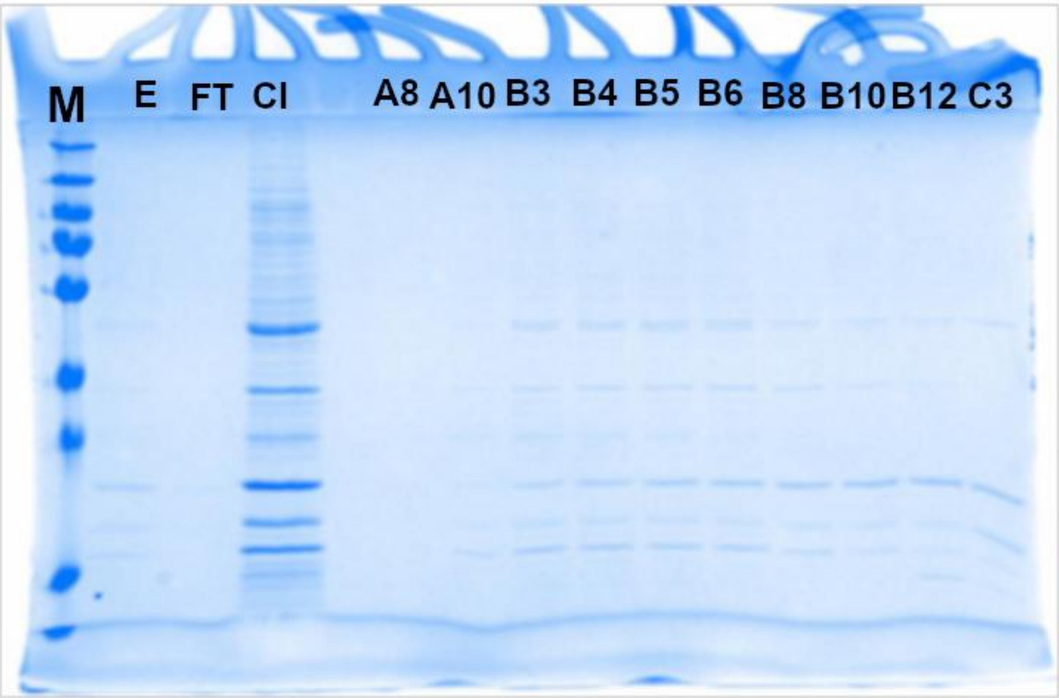


**Supplementary Figure 14**. Full SDS-PAGE of **Fig. 6E** right side gel ('). Mam1ABCD+LucA gel filtration purification fractions of Superose 6 column. The prior step was affinity purification using strep beads, where Mam1D contained a Strep-tag. Fraction B4 shown in **Fig. 6E**. E= Strep-tag elution sample, FT= collected flow through of Strep-tag beads, CI= concentrated elution sample from affinity purification for injection into gel filtration chromatography,Fractions A8 to C3 cover the purified peak from 12 ml to 14.5 ml.

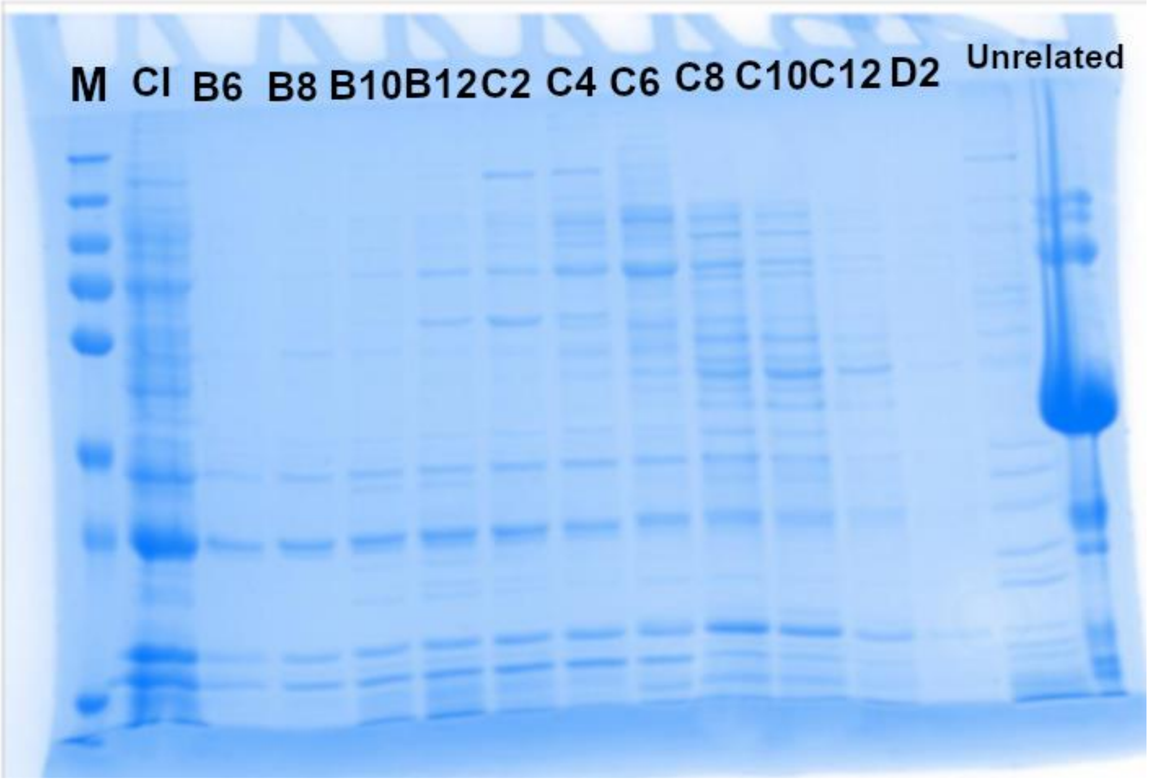


**Supplementary Figure 15**. Full SDS-PAGE figure of **Fig. 6E** left side gel (*). Mam1ABCD Äkta fractions. Mam1D Strep-tag. Fraction B12 lane used in **Fig. 6E**. Fractions cover the elution volume range from 10 ml to 19 ml. Fractions B6 to C4 contain the main peak of the complex, whereas C6 to D2 are assumed to be detached proteins from the main complex, including some impurities. The two rightmost lanes are from unrelated protein purification, not discussed in this paper.

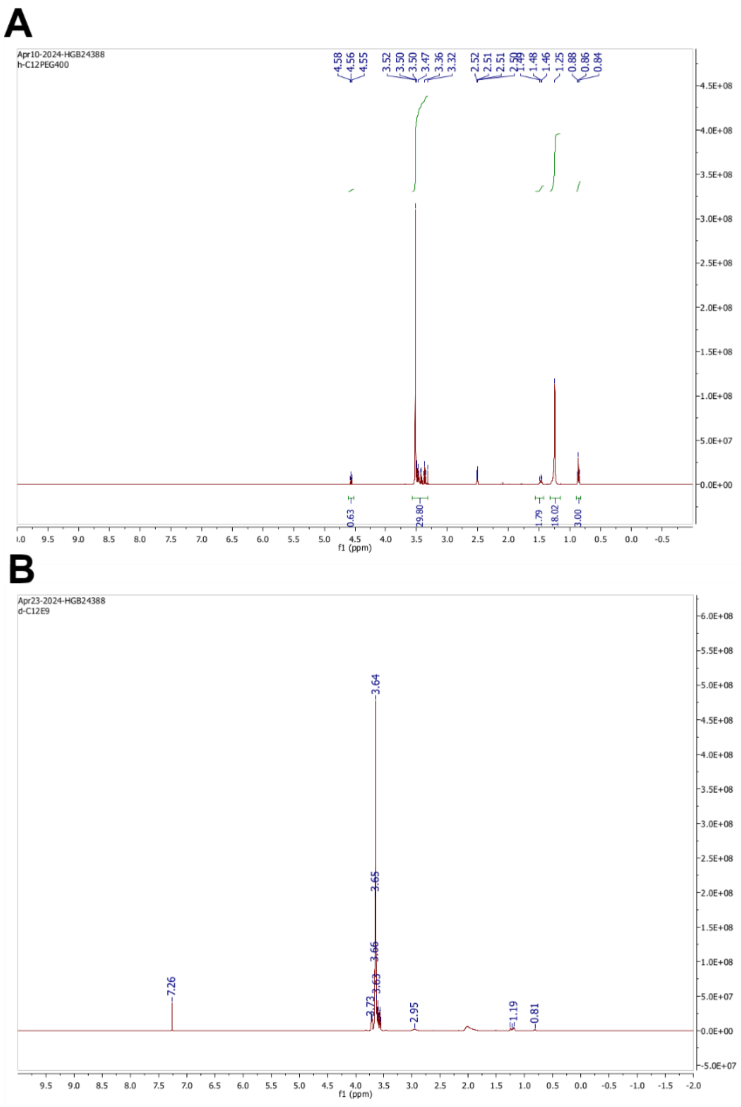


**Supplementary Figure 16. NMR spectrum of d-C12E9**. **(A)** An NMR spectrum of H-C12E9. **(B)** An NMR spectrum of D-C12E9 provided to us by Dr. Kun Ma.

**Supplementary Table 1: Similarities and identities of Mam and Omam proteins with respect to one another**

| | Identity | | | | | | | | | | | | |
|---|---|---|---|---|---|---|---|---|---|---|---|---|---|
| Similarity | M1A | M1B | M1C | M1D | OmA | OmB | OmD | OmC | M3A | M3B | OmE | M4B | M4A |
| Rv0175<br>mam1A | | 4,1 | 21,5 | 18,2 | 23,1 | 15,7 | 25,3 | 22 | 24 | 17,1 | 22,2 | 18,8 | 21,1 |
| **Rv0176<br>mam1B** | 8,3 | | 15 | 11,8 | 6,8 | **15,3** | 13,5 | 10,1 | 10,6 | 11,3 | 10,8 | 12,9 | 14 |
| Rv0177<br>mam1C | 37,7 | 23,3 | | 19,1 | 17,7 | 20,2 | 21,8 | 19,7 | 24 | 21,7 | 20,8 | 15,9 | 16,5 |
| Rv0178<br>mam1D | 34,4 | 22 | 32,4 | | 20,4 | 18,5 | 27,9 | 22,8 | 17,1 | 17,3 | 18,8 | 12,7 | 21,1 |
| Rv0199<br>omamA | 37,6 | 10,5 | 27,6 | 34,3 | | 16,7 | 19 | 20,6 | 24,7 | 20,4 | 16,3 | 9 | 18,1 |
| **Rv0200<br>omamB** | 25,4 | **23,7** | 31 | 27,6 | 26,3 | | 20,2 | 13,7 | 18,1 | 21,5 | 19,4 | 21,9 | 22,2 |
| Rv1362c<br>omamD | 39,4 | 20,3 | 30,7 | 43,5 | 32,3 | 29,6 | | 23 | 21,3 | 31,1 | 24 | 16,7 | 18,8 |
| Rv1363c<br>omamC | 38,6 | 16,2 | 30,9 | 36,4 | 33,8 | 24,8 | 37,6 | | 23,2 | 16,8 | 17,6 | 13,1 | 21,7 |
| Rv1972<br>mam3A | 39,6 | 18,9 | 40,2 | 32,1 | 37,4 | 26,6 | 33,9 | 37,8 | | 20,9 | 24,5 | 15,6 | 18,1 |
| Rv1973<br>mam3B | 27,8 | 17,9 | 31,5 | 27,8 | 32,6 | 32,5 | 42,8 | 26,1 | 35,7 | | 21,9 | 24,4 | 17,7 |
| Rv2390c<br>omamE | 40 | 19 | 33,8 | 35,6 | 35,6 | 33,2 | 40,9 | 32,2 | 41,5 | 39,3 | | 19 | 17,7 |
| Rv3492c<br>mam4B | 30,1 | 19,9 | 32,8 | 26,1 | 13 | 31,6 | 32,8 | 23,6 | 32,7 | 42,7 | 43,6 | | 15,8 |
| Rv3493c<br>mam4A | 32,4 | 23,5 | 31,2 | 32,4 | 29,8 | 13,4 | 29 | 34,3 | 26,6 | 25,6 | 31,9 | 23,9 | |

**Supplementary Table 2: Sequence properties of Mam and Omam genes**

| Protein | TM Helices | TM Aminoacid region | N-terminus facing | N-orientation confidence | Mw, kDa | pI |
|---|---|---|---|---|---|---|
| **Rv0175 Mam1A** | 1 | 55-77 | Cytoplasm | 87% | 22,3 | 4.75 |
| **Rv0176 Mam1B** | 3 | 29-51, 56-78, 166-188 | Cytoplasm | 50% | 35,4 | 11.0 |
| **Rv0177 Mam1C** | 1 | 26-48 | Periplasm | 88% | 25,8 | 5.5 |
| **Rv0178 Mam1D** | 1 | 79-101 | Cytoplasm | 99% | 25,8 | 5.5 |
| **Rv1972 Mam3A** | 1 | 38-60 | Periplasm | 99% | 20,6 | 6.5 |
| **Rv1973 Mam3B** | 1 | 7-26 | Periplasm | 59% | 16,7 | 7.3 |
| **Rv3493c Mam4A** | 1 | 75-97 | Cytoplasm | 45% | 25,5 | 10.4 |
| **Rv3492c Mam4B** | 1 | 5-27 | Cytoplasm | 87% | 18,0 | 9.5 |
| **Rv0199 OmamA** | 1 | 42-64 | Periplasm | 97% | 23,5 | 4.7 |
| **Rv0200 OmamB** | 3 | 7-29, 33-55, 75-97 | Cytoplasm | 99% | 24,0 | 9.3 |
| **Rv1363c OmamC** | 1 | 106-128 | Periplasm | 80% | 28,3 | 4.6 |
| **Rv1362c OmamD** | 1 | 60-82 | Cytoplasm | 62% | 23,5 | 4.4 |
| **Rv2390c OmamE** | 1 | 32-54 | Periplasm | 57% | 19,8 | 9.7 |

N-terminus orientation confidence is taken from TmHMM2.0. Mw and pI are calculated using ProtParam.

**Supplementary Table 3. Protein predictions values of AlphaFold.**

| Protein | pTM | Ranking score | Fraction disordered | Average pLDDT |
|---|---|---|---|---|
| **Mam1A** | 0.62 | 0.79 | 0.33 | 75.32 |
| **Mam1B** | 0.66 | 0.7 | 0.08 | 79.40 |
| **Mam1C** | 0.68 | 0.82 | 0.27 | 82.54 |
| **Mam1D** | 0.64 | 0.85 | 0.41 | 72.15 |
| **Mam3A** | 0.65 | 0.8 | 0.3 | 81.89 |
| **Mam3B** | **0.85** | **0.92** | **0.13** | **92.76** |
| **Mam4A** | 0.61 | 0.81 | 0.41 | 78.21 |
| **Mam4B** | **0.83** | **0.89** | **0.12** | **89.26** |
| **OmamA** | 0.59 | 0.79 | 0.39 | 71.95 |
| **OmamB** | 0.65 | 0.65 | 0.0 | 84.04 |
| **OmamC** | 0.56 | 0.79 | 0.45 | 77.74 |
| **OmamD** | 0.61 | 0.8 | 0.38 | 71.28 |
| **OmamE** | 0.68 | 0.81 | 0.27 | 81.57 |

We have highlighted Mam3B and Mam4B as these two proteins have the shortest variable region sequence, which increases the prediction quality parameters from AlphaFold.
